# Single-Cell Genomics Elucidates Molecular Variations and Regulatory Mechanisms in Circulating Immune Cells

**DOI:** 10.1101/2025.01.26.634963

**Authors:** Jianhua Yin, Yuhui Zheng, Zhuoli Huang, Wenwen Zhou, Yue Yuan, Pengfei Cai, Yong Bai, Shichen Yang, Yue Gao, Shanshan Duan, Yang Wang, Wenxi Zhang, Xinyu Zhang, Yilin Wei, Zekai Xu, Yaling Huang, Ying Liu, Weikai Wang, Tao Yang, Jingzhi Lv, Zhongjin Zhang, Xiaoya Chen, Xiru Zhang, Fupeng Li, Yan Zhang, Guodan Zeng, Xue Wang, Wen Ma, Guixue Hou, Shijie Hao, Chang Liu, Yiwei Lai, Bo Wang, Yuxiang Li, Wenwei Zhang, Peng Gao, Jun Xie, Miguel A. Esteban, Ying Gu, Jiansong Ji, Ting Qi, Boxiang Liu, Jian Wang, Jian Yang, Xun Xu, Longqi Liu, Xin Jin, Chuanyu Liu

## Abstract

The human peripheral blood displays diverse molecular characteristics across populations, understanding the drivers and underlying mechanisms of which remains challenging. Here, we introduce the Chinese Immune Multi-Omics Atlas (CIMA), elucidating sex-, age-, and genetic-related molecular variations by analyzing multi-omics data from 428 adults with over 10 million immune cells. CIMA generated an enhancer-driven gene regulatory network, identifying 237 high-quality regulons and revealing cell type-specific regulatory mechanisms. Additionally, 11,521 lead *cis*-expression quantitative trait loci (eQTLs) and 46,339 chromatin accessibility QTLs (caQTLs) were identified at cell type level. CIMA also uncovered pleiotropic associations among immune-related disease risk loci, eQTLs, and caQTLs in a cell type-specific manner. Lastly, a novel cell language model, CIMA-CLM, was developed to predict chromatin accessibility and noncoding variant effects using chromatin sequences and gene expressions. This work represents a population-scale multi-omics resource of human immune cells, providing a valuable reference for future investigation of immune-related diseases.

## Introduction

The human immune system is a complex cellular network that maintains immune homeostasis, responding to both abnormal physiological and pathological conditions. Molecular variations of immune system arise from a combination of genetic and non-genetic factors (*1–3*), with genetic influences accounting for an estimated 20-40% of inter-individual differences (*4*). A comprehensive understanding of this variation is crucial for elucidating the mechanisms underlying immune-related diseases.

By analyzing the associations between genetic variation and intermediate phenotypes, particularly gene expression, molecular quantitative trait locus (xQTL) studies provide a valuable tool for quantifying the effects of genetic variation on immune system gene expression (*5*). Large-scale projects like BLUEPRINT (*6*), DICE (*7*), and ImmuNexUT (*8*) have employed cell sorting techniques and bulk RNA sequencing (RNA-seq) to the genetic drivers of immune cell transcriptomic variability and to characterize disease risk loci with cell type-specific genetic effects. However, bulk transcriptomic analysis is limited to immune cells identified by known markers and may fail to detect subtle differences in gene expression (*9*). Furthermore, the effect size of single nucleotide polymorphisms (SNPs) often reflects only the average effect in bulk samples, potentially overlooking cellular heterogeneity (*10*).

Recent advancements in single-cell genomics now allow for the study of immune cells at an unprecedented resolution. By integrating single-cell data with genotype information, researchers can link genetic variation to gene expression and chromatin accessibility in immune cells (*11*). As gene expression shifts during immune cell activation and differentiation, single-cell expression QTL (eQTL) analysis enables the identification of genetic influences on these dynamic processes (*12*, *13*). In 2022, the OneK1K project characterized 1.27 million peripheral blood mononuclear cells (PBMCs) from 982 healthy individuals, revealing that a part of genetic variants affect gene expression in a cell type-specific manner (*14*). Similarly, another eQTL study on immune cells from patients with systemic lupus erythematosus (SLE) has annotated genetic variants associated with SLE through fine mapping (*10*). More recently, the Asian Immune Diversity Atlas (*15*) (AIDA) systematically compared the transcriptomic features of immune cells across several East Asian populations, identifying population-specific functional variants through eQTL analysis.

However, a study reported that eQTLs colocalized with a small fraction of immune disease risk loci and suggested that genetic variations more commonly influence the epigenome (*16*). Another study highlighted that disease risk loci can enrich in chromatin regions that are accessible in specific immune cell types (*17*). Shared genetic variation associated with accessible regions and gene expression can occur at transcription factor (TF) binding sites, potentially disrupting enhancer activity and altering gene expression. Therefore, combining analyses of genetic variants, chromatin accessibility, and gene expression can uncover potential causal mechanisms in the regulation of gene expression in immune cells (*18*), significantly improving the interpretation of immune-related disease risk variants. Nonetheless, the limited availability of large-scale single-cell ATAC-seq (scATAC-seq) studies in circulating immune cells has constrained our understanding of these regulatory mechanisms and their roles in diseases.

To address this gap, we present the Chinese Immune Multi-Omics Atlas (CIMA), a systematic analysis of blood components from 428 Chinese adults without active disease. Through independent and integrated multi-omics analyses, we elucidate variations in blood biomolecules and immune cell profiles associated with genetic and non-genetic (sex and age) factors within this population. Using scRNA-seq and scATAC-seq data, we constructed a regulatory network for immune cells. We then characterized the regulatory effects of genetic variation on chromatin accessibility and gene expression in immune cells, revealing cross-omics genetic pleiotropic associations through summary-data-based Mendelian randomization (SMR) analysis (*19*, *20*) with in-house and external genome-wide association studies (GWAS) data (*21*). Furthermore, leveraging our unique scATAC-seq and scRNA-seq data, we developed a novel cell language model, termed CIMA-CLM, which predicts chromatin accessibility and assesses the impact of noncoding variants by integrating chromatin sequences and single-cell gene expression data. Our study not only provides the most comprehensive single-cell multi-omics dataset of the Chinese population to date but also offers new insights into molecular trait variation, laying the groundwork for future mechanistic studies on gene expression regulation in immune cells and the genetic risk of immune-related diseases.

## Results

### A Comprehensive Chinese Immune Multi-Omics Atlas (CIMA) for 428 Individuals

In this study, we performed a comprehensive multi-omics analysis of 428 Chinese adults, including whole-genome sequencing (WGS), scRNA-seq, and scATAC-seq of circulating immune cells, along with biomolecular testing of plasma lipids, metabolites, and blood biochemistry (Fig. 1, A and B and table S1). The cohort consisted of 189 males and 239 females, aged 20 to 77 years, who self-reported no active disease at the time of sampling (Fig. 1B and table S1). After applying quality control measures and removing doublets from PBMCs (fig. S1) (see Methods), we identified 6,484,974 high-quality cells through scRNA-seq (with a median of 15,478 cells per individual) and 3,762,242 high-quality cells through scATAC-seq (with a median of 9,602 cells per individual) (Fig. 1B).

**Fig. 1.**
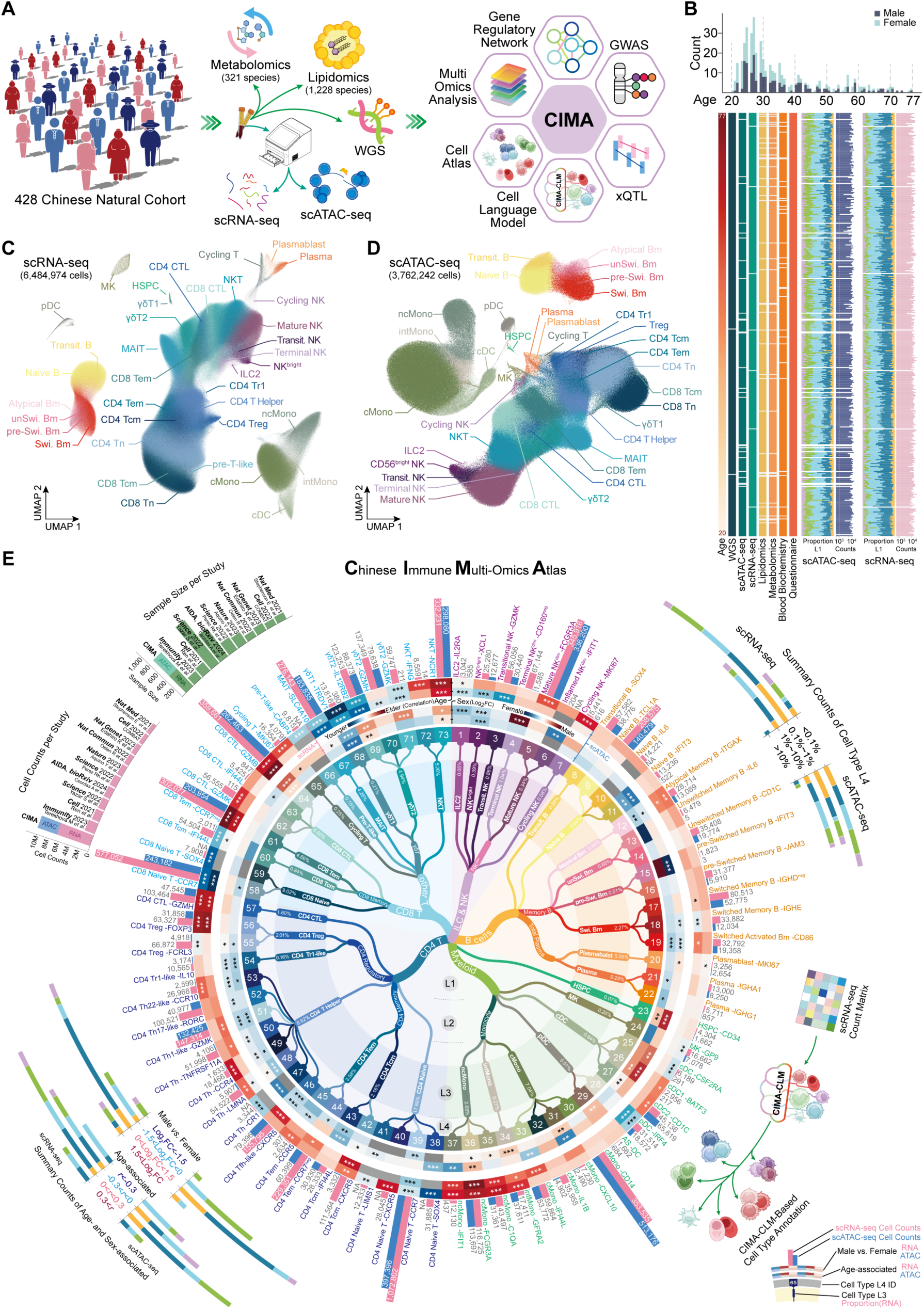
Study design and overview of circulating immune cells for Chinese Immune Multi-Omics Atlas (CIMA). (**A**) Overview of the study design, including collecting peripheral blood samples from 428 Chinese individuals, capturing multi-omics data, and performing comprehensive analysis for CIMA. (**B**) 428 samples arranged in order of increasing age, showing whether different types of data were collected for each sample, the number of cells detected by scRNA-seq and scATAC-seq, and the relative proportions of L1 compartments identified in the study. The top panel shows the distribution of samples by sex and age. (**C**) Uniform manifold approximation and projection (UMAP) plot showing 6,484,974 immune cells from scRNA-seq data that have been subjected to dimension reduction, clustering, and annotation. Cells are categorized into 38 immune cell subtypes at Level 3, with each cell type represented by a different color. (**D**) UMAP plot showing 3,762,242 immune cells from the scATAC-seq data. Annotation labels were transferred from the scRNA-seq data, and each cell type was color-coded (match in **C**). (**E**) A circular dendrogram showing a hierarchy of immune cell annotations based on expression of specific marker genes from scRNA-seq data. L1 represents the five major immune lineages and distinct hematopoietic stem cells (HSPC). L2 (27 subtypes) and L3 (38 subtypes) represent different granularities of cell states. L4 consists of 73 immune cell types marked by specific genes and represented by number and color. The outer ring of the circular dendrogram shows the count of cells captured by scRNA-seq and scATAC-seq for each L4 cell type. The next inner ring shows the correlation between the proportion of each cell type and gender, followed by the ring showing the correlation between the proportion of each cell type and age. A legend is provided in the lower right corner. Bar charts showing the single-cell genomics studies with the highest cell counts in recent years (left top). A bar graph showing the percentage of each L4 cell type among all cells, along with the range of cell percentages, with colors indicating the corresponding L1 lineage (right top). The log2 fold change in the proportions of L4 cells between the sexes, and the range of correlations with age (left bottom). A schematic diagram of the Cell large language (CLM) model for circulating immune cells (right bottom).

To better annotate immune cell phenotypes, we conducted multiple rounds of dimensionality reduction and clustering. Initially, based on the expression of lineage marker genes (fig. S2A), PBMCs were divided into five major cell types at Level 1 (L1): innate lymphoid cells (ILCs) & NK cells, B cells, myeloid cells & HSPCs, CD4 T cells, and CD8 T cells, along with unconventional T cells (see Methods). This Level 1 categorization followed the framework proposed by Silvia Domcke et al. (*22*), with cells representing distinct immune lineages. These five immune lineages were further sub-clustered to identify developmental “cell states” and “cell types” expressing specific genes. Based on classical marker gene expression and cell-type-specific genes (fig. S2, D to H and table S1), we annotated cells at Levels 2 (L2) and 3 (L3) according to their cell states, ultimately identifying 73 immune cell types at Level 4 (L4) with specific functional gene expression (see Methods).

To integrate scRNA-seq and scATAC-seq data, we employed Scglue (*23*), which facilitated the transfer of annotation labels from scRNA-seq to scATAC-seq data. This resulted in high-quality integration (fig. S2C), as reflected by the well-matched cell clusters (Fig. 1, C and D), demonstrating effective data alignment. The hierarchical structure of the annotated immune cells is shown (Fig. 1E). Rare cell types, such as AS-DC cells, which exhibit characteristics of both cDC and pDC cells (*24*), were identified at Level 4, with proportions below 0.1% in both scATAC-seq and scRNA-seq datasets (Fig. 1E, right top and table S1). Additionally, we identified cell types associated with sex and age (Fig. 1E, left bottom). For further investigation of immune effects specific to cell types, we selected L4 cells for subsequent analyses. Using these L4 cells as a reference, we developed a model for the fine annotation of human PBMC subtypes (Fig. 1E, right bottom).

In comparison to the top 10 studies with the highest single-cell counts in PBMC cohort research in recent years, which all employed single-cell transcriptomics, our study stands out by incorporating both scRNA-seq and scATAC-seq, with a combined cell count exceeding 10 million (Fig. 1E, left top). Thus, our study provides an extensive and highly detailed multi-omics dataset from the Chinese population (CIMA), enabling high-resolution identification of circulating immune cells. Furthermore, it offers a robust model for the annotation of human PBMCs, which can serve as a valuable resource for future research.

### Biomolecular and immunological characteristics in blood exhibit sex- and age-related variations

CIMA offers a unique opportunity to investigate the diversity and variation in blood multi-omics molecular characteristics within the Chinese population. We began by performing principal component analysis (PCA) on five types of individual-level bulk data, which included blood biomolecules and immune cell proportions derived from scRNA-seq and scATAC-seq data (Fig. 2A). The first two principal components (PC1 and PC2) of the cell proportions clearly distinguished age groups, while metabolic and biochemical molecules showed sex-related differences in their principal components (Fig. 2B). Then we used Multi-Omics Factor Analysis (MOFA) to integrate multiple data modalities and identifies latent factors that capture the multi-omics inter-individual variability. The biological interpretation of latent factors can be provided by the factor-related high-weight features (*25*). Sample-level data from 428 individuals, including 19 blood biochemical markers, 1,228 lipids, 321 metabolites, and proportions of 73 cell types identified by scRNA-seq and 65 cell types identified by scATAC-seq were input and we identified a sex associated factor (Fig. 2, A and C and fig. S3A). The factor included levels of 2,3-diphosphoglyceric acid and the proportion of CD8 CTL-GZMB cells, which were significantly elevated in males (Fig. 2D). After correcting for sex-related effects, we extracted a set of age-related multi-omics components and found that certain phospholipids and immune cell type proportions carried high feature weights in Factor 2 (Fig. S3B and table S2). Previous studies have shown that phospholipid accumulation occurs in a sex-conserved manner during aging (*26*). Additionally, an increase in ncMono-FCGR3A cells and CD4 CTL-GZMH cells were observed (Fig. 2E). The expansion of CD4 CTL-GZMH cells with age has been linked to the maintenance of healthy immune cytotoxic functions in centenarians (*27*).

**Fig. 2.**
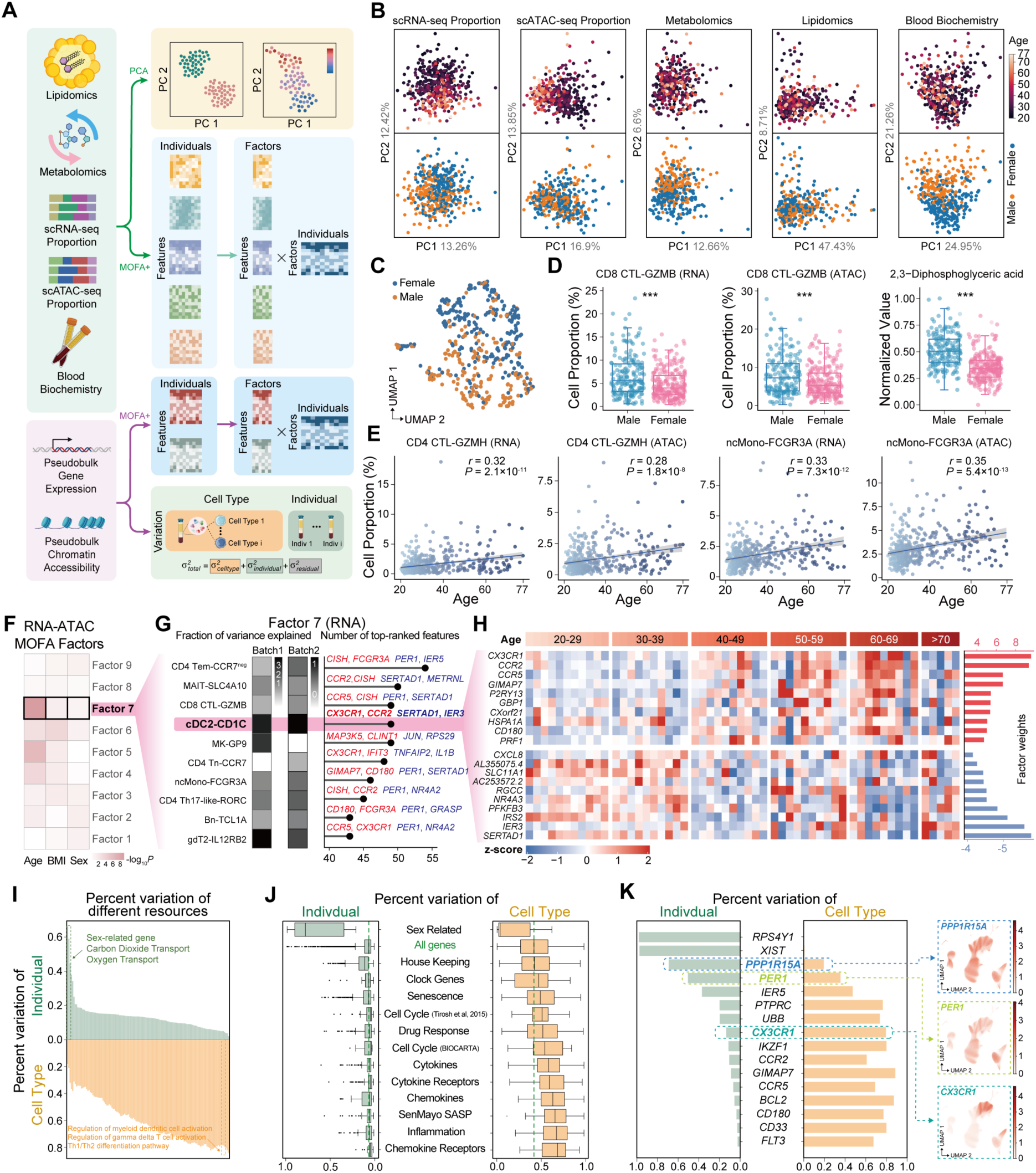
Age- and sex-related multi-omics molecular characteristics. (**A**) Overview of the workflow for multi-omics analysis and the datasets analyzed. (**B**) PCA results of multi-omics datasets, including cell type proportions from scRNA-seq and scATAC-seq, as well as data from metabolomics, lipidomics, and blood biochemistry. (**C**) UMAP plot showing the distribution of sexes (female and male) in the integrated MOFA model using multi-omics datasets, including cell type proportions from scRNA-seq and scATAC-seq, metabolomics, lipidomics, and blood biochemistry. (**D**) Box plot illustrating sex differences (female and male) in the proportions of CD8 CTL-GZMB cells (RNA), CD8 CTL-GZMB cells (ATAC), and 2,3-Diphosphoglyceric acid. (**E**) Linear regression analysis estimating the relationship between age (x-axis) and cell type proportion (y-axis) for CD4 CTL-GZMH cells (RNA), CD4 CTL-GZMH cells (ATAC), ncMono-FCGR3A cells (RNA), and ncMono-FCGR3A cells (ATAC). Each dot represents one individual. (**F**) Correlation analysis of the top 9 integrated MOFA factors with clinical indicators using pseudo-bulk ATAC-seq and RNA-seq datasets. (**G**) Heatmap showing the percentage of variance explained by each of the top 10 views (cell types) for factor 7 (RNA) in batch 1 and batch 2, respectively (left). Bar plots showing the number of top-ranked features for each of the top 10 views (right). (**H**) Normalized expression values of selected top-ranked features across different age stages in cDC2-CD1C cells (left). Bar plot showing the feature factor weight (right), with plus and minus signs indicating the direction. (**I**) Bar plots showing the percentage of variation attributed to individual (green) and cell type (yellow) for collected Gene Ontology (GO) functions, with selected GO categories highlighted. (**J**) Box plots showing the distribution of percentage variation attributed to individual (green) and cell type (yellow) for specific gene sets, with all genes as the control gene set. (**K**) Bar plots showing the percentage of variation attributed to individual (green) and cell type (yellow) for specific genes, with selected genes highlighted (left). UMAP plot of example genes, with *PPP1R15A* and *PER1* demonstrating cell type-shared expression patterns contributing to high individual variability, and *CX3CR1* demonstrating cell type-specific expression patterns contributing to high cell type variability (right).

MOFA analysis were also performed on gene expression and chromatin accessibility data at the immune cell type level (Fig. 2A), and identified Factor 7 as the most age-associated factor (Fig. 2F). Factor 7 explained most of the variance in cDC2-CD1C cells (Fig. 2G and table S2). By examining the high-weight features in cDC2-CD1C cells, we observed age-related increases in the expression of *CX3CR1* and *CCR2*, and decreases in the expression of *SERTAD1* and *IER3* (Fig. 2, G and H). Gene Ontology (GO) function enrichment analysis indicated that the genes upregulated with age were involved in processes such as “cellular defense response”, “positive regulation of response to external stimulus”, and “response to bacterium” (fig. S3C), suggesting that immune responses to external stimuli may accumulate with age, as reflected by gene expression changes. We next examined the characteristics of high-weight genes (Factor weight > 3 or Factor weight < −3) that are either upregulated (positive weight) or downregulated (negative weight) with age (fig. S3D). We found that a significant proportion of both age-upregulated (58%) and age-downregulated (46.4%) genes are cell type-specific (fig. S3E). These distinct gene patterns were enriched in different functional pathways, shedding light on the biological processes driving these age-related changes (fig. S3F).

To further investigate the sources of gene expression variations, we applied the variancePartition (*28*) algorithm to perform variance decomposition analysis (Fig. 2A). Pathways with higher cell type variation were associated with immune activation, differentiation, and regulatory functions, such as the “Th1/Th2 differentiation pathway”. In contrast, pathways showing higher individual variation were linked to essential physiological processes, such as “oxygen transport” and “carbon dioxide transport” (Fig. 2I and table S2). As anticipated, genes related to sex exhibited the highest degree of individual variation (*29*) (Fig. 2, I and J). We obtained similar results in calculating the percent variance of interested gene sets (Fig. 2, J and K). Notably, *PER1* and *PPP1R15A* displayed substantial individual differences, were ubiquitously expressed across cell types, and showed decreased expression with age (Fig. 2K and fig. S2G). Both *PER1* and *PPP1R15A* play roles in the regulation of aging (*30*, *31*). In contrast, *CX3CR1* exhibited high variation across cell types and was specifically expressed in NK cells (Fig. 2K). This analysis comprehensively considered both cell type and individual sources of variation, suggesting that aging-related gene expression is not only finely regulated within specific cell types but also influenced across multiple cell types over time.

In summary, MOFA provided a comprehensive collection of multi-omics features related to sex and age, offering a valuable resource for exploring the cellular and molecular mechanisms regulating gene expression in immune cells.

### Construction of immune cell type-specific gene regulatory networks

To identify candidate *cis*-regulatory elements (cCREs) in each immune cell types, we performed peak calling of chromatin accessibility signals using MACS3 within the SnapATAC2 workflow (*32*). This analysis resulted in the identification of 338,036 open chromatin regions (each extending 501 bp around the peak summit), referred to as cCREs (table S3). These peaks were annotated using HOMER (*33*). Our findings showed that only 6.9% of the cCREs were located in promoter regions of protein-coding and long noncoding RNA genes. In contrast, 31.2% were found in intronic regions, 20.7% in intergenic regions, and 32.9% in transposable elements (TEs), including long terminal repeats (LTRs), long interspersed nuclear elements (LINEs), short interspersed nuclear elements (SINEs), RNA repeats, DNA repeats, and other repeats (Fig. 3A). Notably, our identified cCREs substantially expand upon previously defined human cCREs based on bulk chromatin data from the ENCODE project (*34*). Specifically, 57.14% of the circulating immune cell cCREs (table S3) did not overlap with DNase-hypersensitive sites (DHSs) mapped across various human tissues (Fig. 3B, left). Further analysis revealed that 51.5% of the overlapping cCREs were active in more than ten cell types, while only 17.3% were accessible in a single cell type. In contrast, over 51% of the non-overlapping cCREs were accessible in only one cell type, with less than 10% being active in more than ten cell types (Fig. 3B, right).

**Fig. 3.**
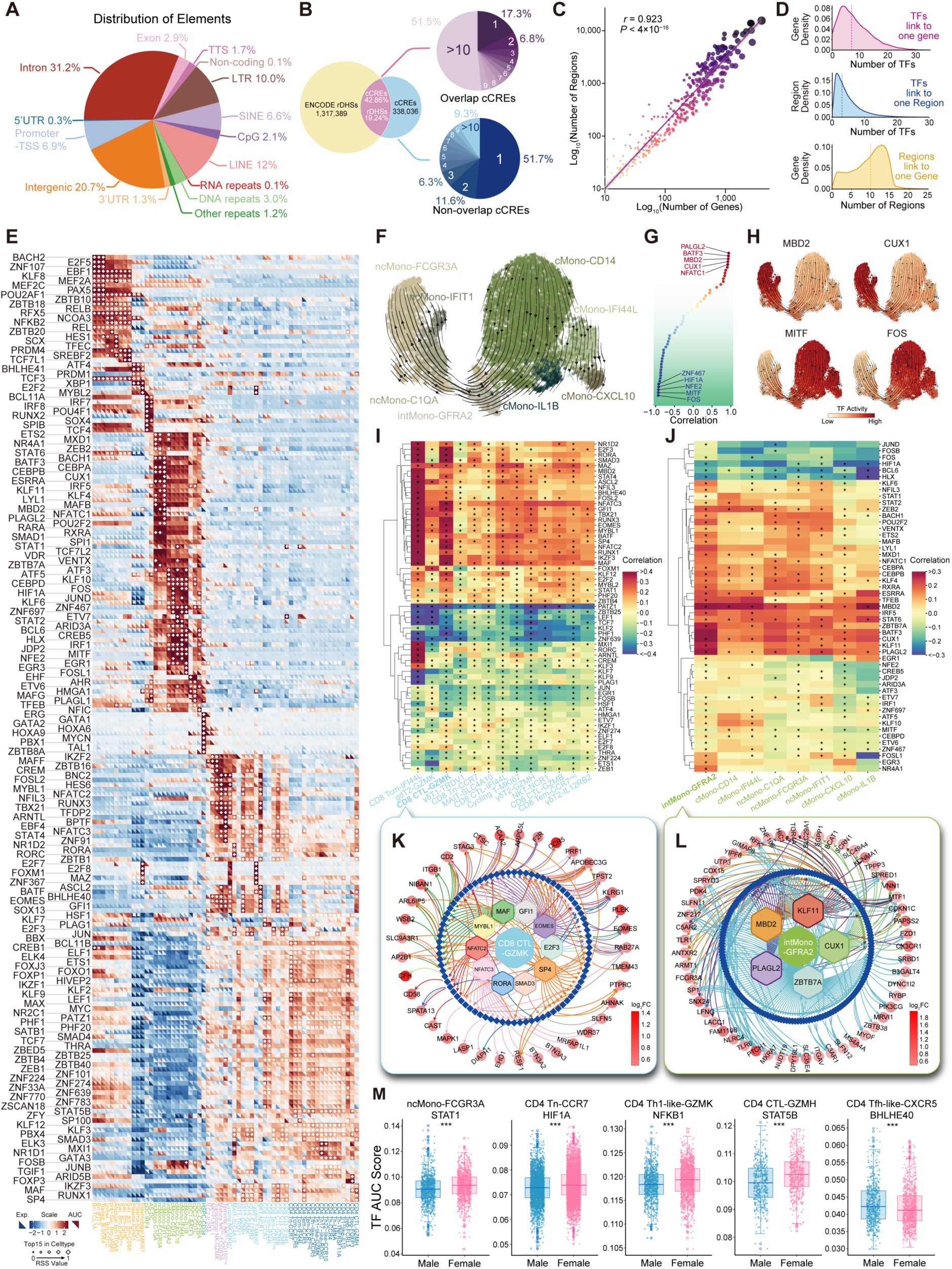
Construction of immune cell-specific gene regulatory networks. **(A)** Annotation and proportion of cCREs. (**B**) Overlap of cCREs in this study with ENCODE rDHSs. Statistical distribution of cCREs overlapping with rDHSs across different cell types (right top), and not overlapping with rDHSs across different cell types (right bottom). (**C**) Correlation analysis between the number of genes and regions regulated by each TF. (**D**) Distribution of the number of TFs linked to each gene (top), the number of TFs linked to each region (middle), and the number of regions linked to each gene (bottom). (**E**) Heatmap of TF expression levels and AUC scores, and dot plots showing the cell type-specific scores (RSS scores) of the top 15 transcription factors in each cell type. The plot uses a color scale to display TF expression levels and AUC scores, and a dot size scale to display RSS values. (**F**) Inferring the differentiation trajectory of monocytes through scTour. (**G**) Correlation between TF activity and differentiation pseudotime in monocytes. (**H**) Dynamics of TF activity during cell differentiation in monocytes. (**I and J**) Heatmap showing correlation between transcription factor activity and age in CD8 T cells & unconventional T cells (I) and monocytes (J). (**K and L**) Age-related GRNs in CD8 CTL-GZMK (K) and intMono-GFRA2 (L) were constructed based on transcription factors whose activity is upregulated with aging, regions that accessible within these cell types, and target genes whose expression is upregulated with aging. (**M**) Violin plot showing transcription factors with significant activity differences between males and females.

CREs modulate gene expression by interacting with sequence-specific TFs (*35*). By integrating scRNA-seq and scATAC-seq data, we constructed an enhancer-driven gene regulatory network (GRN) within the L4 cell type (*36*). After filtering out regions with negative region-to-gene correlations, we identified 404 eRegulons, encompassing 84,625 targeted regions and 13,645 targeted genes (table S4). A positive correlation was observed between the number of genes and regions regulated by each TF (Fig. 3C). Our analysis revealed that 97% of genes were associated with 1 to 15 predicted enhancers, while 95% were regulated by 1 to 20 TFs (Fig. 3D).

Given the highly detailed annotations we provided at L4 cell types, GRNs are essential for gaining a deeper understanding of cellular identities. Dimensionality reduction based on eRegulon enrichment scores effectively distinguished between different cell types (fig. S4A). We selected 237 high-quality eRegulons and identified TFs specific to each cell type (Fig. 3E, fig. S4, B and C and table S4). This GRN highlighted numerous known TFs involved in specific immune cell differentiation, such as PAX5, BACH2, EBF1, PRDM1, and XBP1 for B cell differentiation (*37*); SPI1, CEBPB, CEBPA, and NR4A1 for monocyte regulation (*38*); and RUNX3, STAT4, GATA3, and EOMES for T cell differentiation (*39*). Additionally, TFs specific to rare cell types were identified. In cMono-IL1B cells, the most specific TFs were ATF3, BCL6, FOSL1, EGR1, and EGR3, with ATF3 regulating the most differentially expressed genes upregulated in this cell type (*CXCL3*, *CCL3*, *CXCL8*, *NLRP3*) (fig. S4D). In cMono-CXCL10 cells, the most specific TFs were ATF3, STAT1, STAT2, ETV7, and JDP2 (fig. S4D), with STAT1 regulating the most differentially expressed genes upregulated in this cell type (*CXCL10*, *CCL2*, *TNFSF10*, *TNFSF13B*) (fig. S4E). Using hierarchical clustering, we grouped all TFs into seven distinct modules (fig. S4F). These modules are associated with key processes such as T cell differentiation and activation, B cell differentiation, humoral immune response, and myeloid cell differentiation, highlighting the critical roles of TFs in regulating immune cell differentiation. We further examined the dynamic activity of TFs during monocyte differentiation (Fig. 3, F and G). During the transition from classical monocytes to non-classical monocytes, we observed notable changes in the activity of TFs such as MITF, FOS, MBD2, and CUX1 (Fig. 3H). Accordingly, we identified functional TFs that may encode cell identity and involve in cell differentiation, which could serve as additional useful signatures for characterizing immune cell types (table S4).

To determine whether TF activity varies among individuals, we assessed the correlation between cell type-specific GRNs and aging (Fig. 3, I and J). In CD8 T and unconventional T cells, we identified several TFs significantly correlated with aging, including EOMES, RUNX3, and TBX21, which are essential for regulating T cell function (*39*). We then constructed an age-related GRN, focusing on TFs and their target genes that were significantly upregulated with age. Among the TFs with increased activity during aging, NFATC3 regulated the most age-related genes, such as *PRF1*, *SLFN5*, *BTN3A2*, and *BTN3A3* (Fig. 3K). EOMES was linked to changes in genes like *CCR5* and *F2R*. In monocytes, TFs such as MBD2, CUX1, and ZBTB7A showed significant upregulation in activity with age. Specifically, in intMono-GFRA2 cells, ZBTB7A regulated the most genes upregulated during aging, including *C3AR1*, *TLR1*, *TLR6*, and *FCGR3A*; MBD2 regulated *C5AR2* and *TLR1*, while CUX1 regulated *CX3CR1* (Fig. 3L). In addition to age-related findings, our GRN analysis revealed TFs with significant activity differences between males and females (Fig. 3M and table S5). In CD4 T cells with notable sex-based differences in cell proportions, we observed that HIF1A, NFKB1, and STAT5B had higher activity in females, whereas BHLHE40 showed higher activity in males. In ncMono-FCGR3A cells, STAT1 activity was higher in females.

In summary, we have established GRNs specific to immune cell types, identified a range of TFs that linked to immune cell differentiation, aging, and sex. These findings have significantly deepened our comprehension of the sources of immune diversity.

### Identification of eQTLs and caQTLs at the level of immune cell types

Chromatin accessibility and gene expression are influenced by genetic variation within populations, which may reflect underlying regulatory mechanisms. To investigate the relationship between genetic variation, chromatin accessibility, and gene expression, we generated genotype data by performing WGS of the CIMA cohort. Variants were imputed using the high-coverage 1000 Genomes Project (1kGP) imputation panel, which includes 3,202 samples (*40*). After quality control, 8,766,136 variants with a minor allele frequency > 0.01 were retained for xQTL analysis (see Methods). PCA analysis of these variants demonstrated that individuals from the CIMA cohort are closely clustered with individuals of East Asian ancestry from the 1kGP (fig. S5A).

To map *cis*-eQTLs and *cis*-caQTLs, we created pseudobulk measures of RNA expression and chromatin accessibility for each cell type. This was achieved by aggregating normalized RNA expression or chromatin accessibility from all cells assigned to a given cell type within each individual (see Methods). After removing cell types with fewer than 70 pseudobulk samples, we proceeded with the analysis at Level 4, covering 69 cell types for *cis*-eQTL mapping and 41 cell types for *cis*-caQTL mapping. The TensorQTL was used to identify xQTLs within a ±1 Mb region of the gene’s transcription start site (TSS) or the midpoint of chromatin accessibility peaks. Age, sex, genotype principal components (PCs), and probabilistic estimation of expression residuals (PEER) factors were included as covariates (see Methods). The summarized *cis*-xQTL mapping results for L4 cell types are provided (table S6 and S7). We summarized the number of eGenes and caPeaks detected in each cell type (Fig. 4, A and B). An “eGene” refers to a gene where the variant-gene QTL pair exceeds the significance threshold in TensorQTL, using a two-step false discovery rate (FDR) of < 0.05 (see Methods). Similarly, a “caPeak” represents a chromatin accessibility peak where the variant-gene QTL pair meets the same significance threshold. In total, we identified 11,521 eGenes, of which 3,022 (26%) were detected in only one cell type (Fig. 4A), and 46,399 caPeaks, of which 28,792 (62%) were cell type-specific (Fig. 4B). These results indicate the presence of cell type-specific gene regulation. The substantial variability in the number of eGenes and caPeaks across cell types can be partially explained by differences in cell type proportions, as we observed a significant positive correlation between these counts and cell type proportions (Fig. 4, C and D), consistent with previous studies (*41*).

**Fig. 4.**
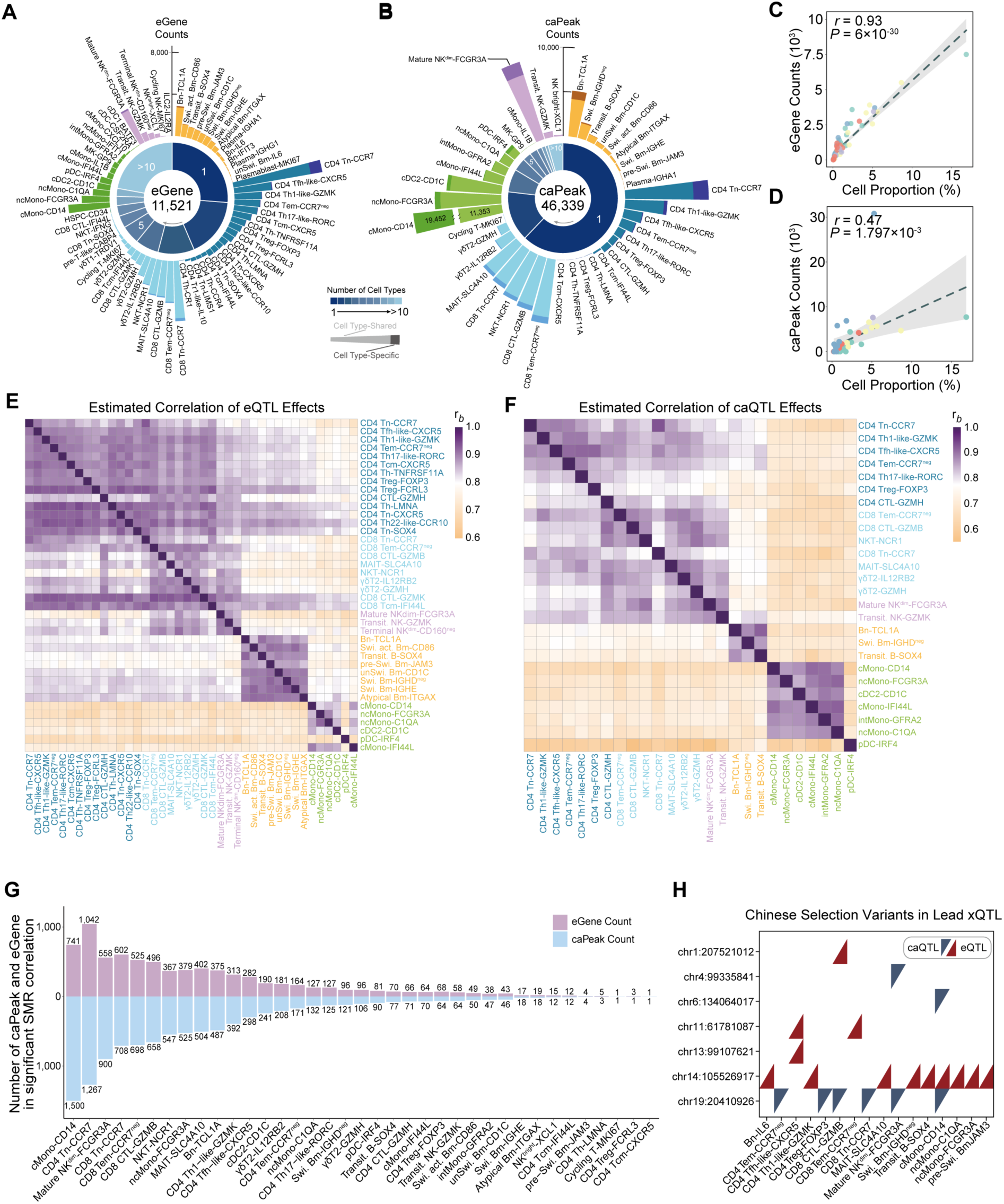
Summary of eQTL and caQTL in immune cell types. **(A)** Bar plot illustrating the number of eGenes detected across the 69 cell types included in the eQTL analysis. The pie chart displays the distribution of significantly detected eGenes, categorized by the number of cell types in which they were identified. **(B)** Bar plot illustrating the number of caPeaks detected across the 41 cell types included in the caQTL analysis. The pie chart displays the distribution of significantly detected caPeaks, categorized by the number of cell types in which they were identified. (**C and D**) Correlation between cell proportions and the number of eGenes (C) and caPeaks (D). (**E**) Heatmap displaying the estimated correlations of eQTL effects (*r_b_*) between cell types with more than 500 eGenes. Rows represent reference eQTLs, while columns represent query eQTLs. (**F**) Heatmap displaying the estimated correlations of caQTL effects (*r_b_*) between cell types with more than 500 eGenes. Rows represent reference caQTLs, while columns represent query caQTLs. (**G**) Bar plot illustrating the number of eGenes and caPeaks detected in significant variant-caPeak-eGene SMR pleiotropic associations across cell types. (**H**) Overlapping loci between lead variants of eQTLs/caQTLs and the 98 loci under selection across the latitudinal gradient in China.

To validate the robustness of our eQTL findings, we compared them with datasets from populations of different ancestries. Among the 11,521 eGenes, 4,626 (40.15%) overlapped with the OneK1K (*42*) dataset (European ancestry), and 10,634 (92.30%) overlapped with the ImmuNexUT (*8*) dataset (East Asian ancestry) (fig. S5, B and C). Both OneK1K and ImmuNexUT provide *cis*-eQTLs for various immune cell types. We further analyzed eQTL sharing across cell types using the *π*1 statistic (see Methods). The eQTLs in various cell types from OneK1K and ImmuNexUT were broadly shared with our findings (fig. S5, D and E). Additionally, we observed distinct stratification in the sharing of results across different cell types in our data compared to those from ImmuNexUT. Since ImmuNexUT cell types are derived from bulk samples through flow cytometry sorting, our results suggest that single-cell omics clustering may capture more cell type-specific patterns with higher resolution (fig. S5E). Furthermore, we have provided the most comprehensive caQTL dataset available for immune cells.

Next, we applied the *r_b_* method to detect associations of xQTL genetic effects across different cell types (see Methods). Our analysis revealed that cell types within the same lineage exhibited stronger associations in their genetic effects. For instance, xQTL genetic effects were similar among T cells (Fig. 4, E and F), while plasmacytoid dendritic cells (pDCs) displayed more distinct xQTL genetic effects (Fig. 4, E and F).

To investigate whether the effect of a genetic variant on gene expression is mediated by chromatin accessibility, we employed SMR on caQTL and eQTL results to test for pleiotropic associations between chromatin accessibility and gene expression in each overlapping cell type (*19*) (see Methods). We identified 6,930 peak-gene pairs with significant SMR associations, involving 4,979 caPeaks and 2,602 eGenes across 37 cell types. Among these, cMono-CD14 cells exhibited the highest number of significant variant-peak-gene SMR relationships, with 1,500 caPeaks and 741 eGenes (Fig. 4G). Given the CIMA cohort’s broad geographic coverage, spanning from northern to southern China, genetic variation may be influenced by regional ancestry differences. As such, some xQTL effects could reflect differences between northern and southern Chinese populations. To explore this, we examined 98 previously reported loci associated with latitude-based selection across China (*43–47*). We found that seven lead xQTLs in our study overlapped with these loci, involving 16 cell types. These findings provide insights into the potential genetic effects of these loci on L4 cell types (Fig. 4H and table S8).

In summary, we provide a comprehensive resource of cell type-specific eQTLs and caQTLs for immune cells, using genetic variation as instrumental variables to study potential regulatory relationships between caPeaks and eGenes. Additionally, we annotated loci influenced by selection in northern and southern Chinese populations using xQTL data.

### Genetic pleiotropic associations within multi-omics data indicate the effects of selection loci in Chinese populations

We further utilized our cell type-specific xQTL results to explore potential associations between caPeaks, eGenes, and traits, with a focus on cell type-specific patterns. Specifically, we applied the SMR approach to each xQTL result at the cell type level to assess whether a caPeak or eGene is associated with a trait and to determine if the caPeak-trait or eGene-trait association is driven by the same causal variants (Fig. 5A) (see Methods).

**Fig. 5.**
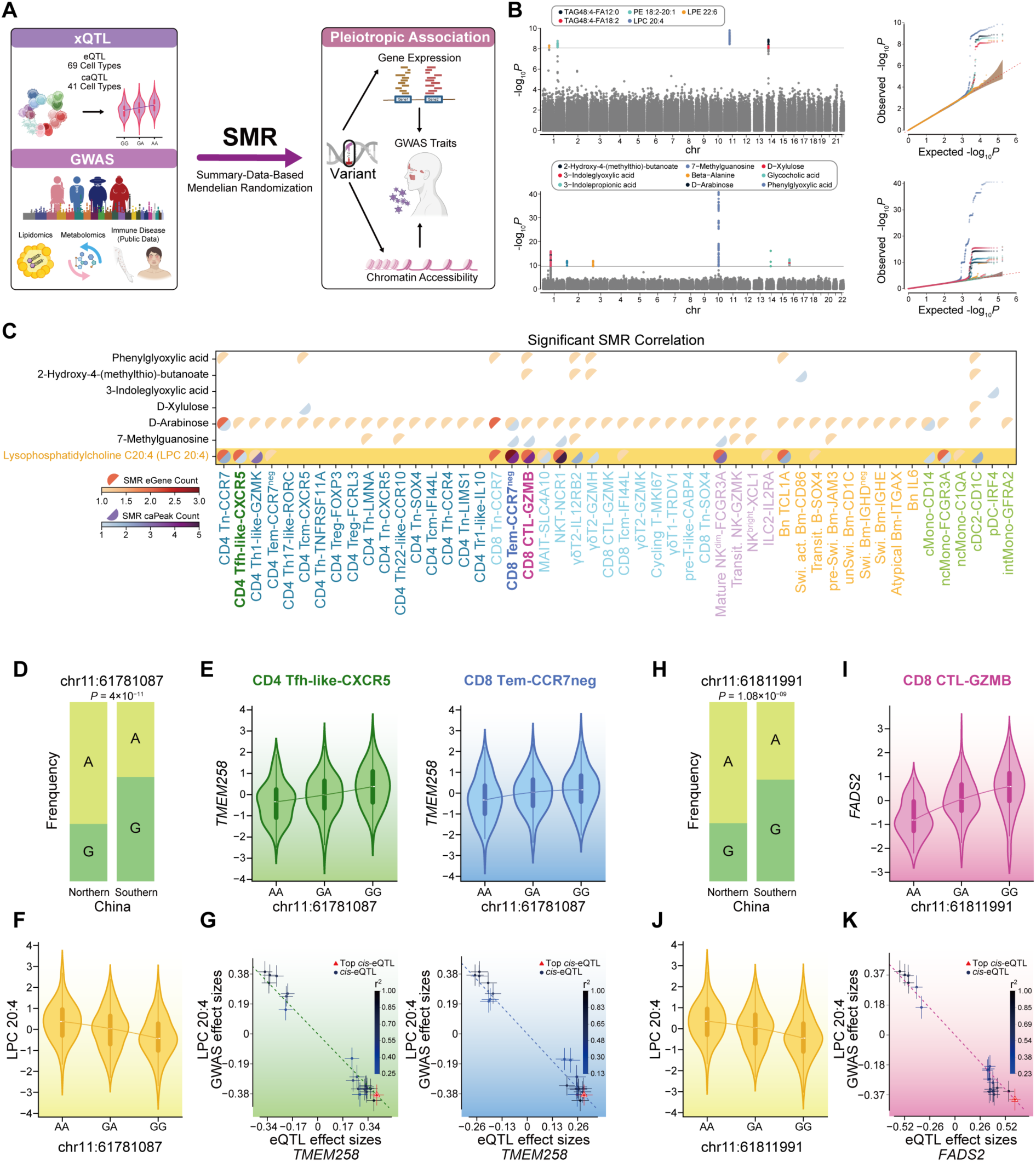
Genetic pleiotropic associations among xQTL, lipidome, and metabolome. (**A**) Workflow of SMR analysis using xQTL and GWAS summary statistics. (**B**) Manhattan plots and QQ plots showing significant associations with 5 lipids and 9 metabolites. (**C**) Heatmap illustrating the number of eGenes and caPeaks detected in significant variant-caPeak/eGene-lipid/metabolite SMR pleiotropic associations across cell types. (**D**) Bar plot illustrating the differences in allele frequency at the chr11:61781087 between northern and southern China. (**E**) Violin plot illustrating the differences in *TMEM258* expression of CD4 Tfh-like-CXCR5 cells (left) and CD8 Tem-CCR7^neg^ (right) across various genotypes at the chr11:61781087. (**F**) Violin plot illustrating the differences in LPC 20:4 level across various genotypes at the chr11:61781087. (**G**) Effect sizes of variants from LPC 20:4 GWAS plotted against those for variants from the *TMEM258* eQTL in CD4 Tfh-like-CXCR5 cells (left) and CD8 Tem-CCR7^neg^ (right). The orange dashed lines represent the estimate of *b*_xy_ at the top *cis*-eQTL. Error bars are the standard errors of variant effects. (**H**) Bar plot illustrating the differences in allele frequency at the chr11:61811991 between northern and southern China. (**I and J**) Violin plot illustrating the differences in *FADS2* expression of CD8 CTL-GZMB cells across various genotypes at the chr11:6181191 (I), and LPC 20:4 level across various genotypes at the chr11:6181191 (J). (**K**) Effect sizes of variants from LPC 20:4 GWAS plotted against those for variants from the *FADS2* eQTL in CD8 CTL-GZMB cells. The orange dashed lines represent the estimate of *b*_xy_ at the top *cis*-eQTL. Error bars are the standard errors of variant effects.

The traits of interest included lipidomics, metabolomics, and immune-related diseases. To obtain GWAS summary statistics for these traits, we conducted in-house GWAS analyses based on our lipidomics and metabolomics data (see Methods). Additionally, we gathered GWAS summary statistics for 52 immune-related diseases from public datasets (table S9), with a focus on individuals of East Asian ancestry (*21*). Through GWAS of 623 lipids and 321 metabolites, we identified significant associations with five lipids at 133 variants and nine metabolites at 343 variants (Fig. 5B and table S10).

Among these 14 lipids and metabolites, the SMR method detected 136 significant pleiotropic associations across 50 cell types, involving 11 eGenes, 12 caPeaks, 1 lipid, and 6 metabolites (Fig. 5C). One example is lysophosphatidylcholine (LPC) 20:4. Two variants, chr11:6171087 and chr11:61811991, which display selective effects in northern and southern China, were pleiotropically associated with gene expression in specific cell types and plasma levels of LPC 20:4 (table S11). The allele A of variant chr11:6171087 is more frequent in northern China (Fig. 5D). As the genotype shifts from AA to GG, the expression levels of *TMEM258* in CD4 Tfh-like-CXCR5 and CD8 Tem-CCR7^neg^ cells increase (Fig. 5E), while plasma levels of LPC 20:4 decrease (Fig. 5F), indicating an inverse correlation between the variant’s effect on LPC 20:4 and its effect on *TMEM258* (Fig. 5G). This finding suggests that gene expression changes related to protein folding and endoplasmic reticulum stress in CD4 Tfh-like-CXCR5 and CD8 Tem-CCR7^neg^ cells may influence fatty acid metabolism.

Similarly, we identified another locus, variant chr11:6181191, where allele A is more frequent in northern China (Fig. 5H). As the genotype changes from AA to GG, the expression of *FADS2* in CD8 CTL-GZMB cells increases (Fig. 5I), while plasma levels of LPC 20:4 decrease (Fig. 5J), suggesting that the variant’s effect on LPC 20:4 is inversely correlated with its effect on *FADS2* (Fig. 5K). Fatty acid desaturase 2 (encoded by *FADS2*) is a key enzyme in arachidonic acid biosynthesis, and LPC 20:4 contains a 20:4 fatty acid chain, which can be converted into arachidonic acid through multiple enzymatic reactions (*48*). Thus, one possible hypothesis is that increased *FADS2* expression in CD8 CTL-GZMB cells enhances desaturase activity, increasing arachidonic acid synthesis, thereby reducing the demand for LPC 20:4, resulting in a negative feedback mechanism that decreases LPC 20:4 levels.

Therefore, we propose that selection loci in the Chinese population may mediate changes in plasma lipid and metabolite phenotypes by modulating gene expression in specific cell types.

### Identification of cell type-specific multi-omics immune mediators of disease risk loci

Immune cells play a crucial role in the development and progression of various diseases, with disease-associated risk loci potentially modulating chromatin accessibility and gene expression in a cell type-specific manner. By applying the SMR method using GWAS summary statistics from 52 immune-related diseases to our xQTL results, we identified 1,727 significant pleiotropic associations across 69 cell types, involving 113 eGenes, 195 caPeaks, and 21 diseases (Fig. 6A and table S12). Notably, 17 variants showed pleiotropic associations with caPeaks, eGenes, and diseases simultaneously.

**Fig. 6.**
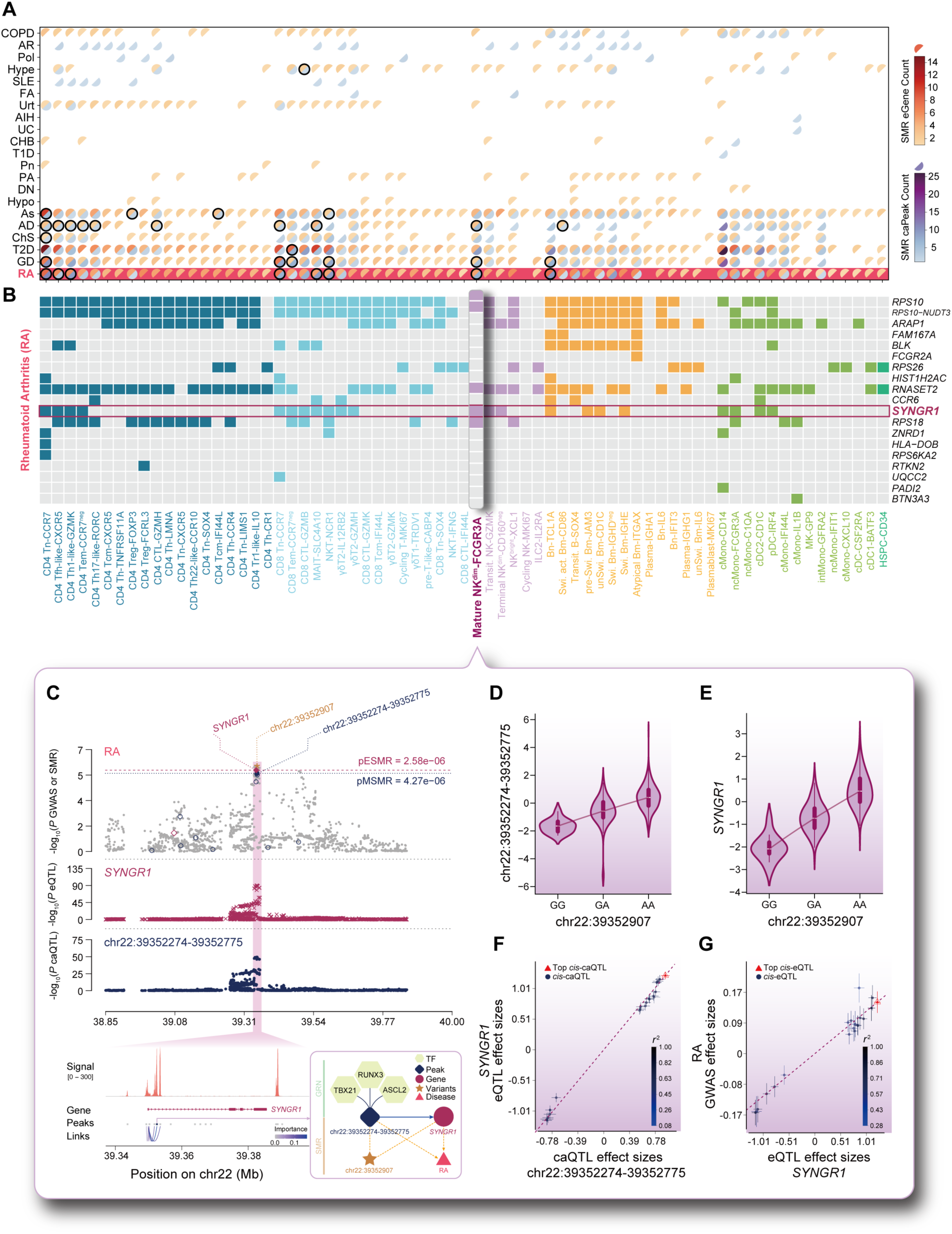
Genetic pleiotropic associations between xQTL and immune related disease loci. **(A)** Heatmap illustrating the number of eGenes and caPeaks detected in significant variant-caPeak/eGene-disease SMR pleiotropic associations across cell types. The black outline indicates that the shared variant simultaneously pleiotropically associated with caPeak, eGene, and disease was detected. (**B**) The eGenes detected in significant variant-eGene-disease SMR pleiotropic associations across cell types in RA. (**C**) Results of variants and SMR associations across caQTL, eQTL, and RA GWAS in Mature NK^dim^-FCGR3A cells. The top plot shows −log_10_(*P* values) of variants from the GWAS of RA. The red diamonds and blue circles represent −log_10_(*P* values) from the SMR tests for associations of eGenes and caPeaks with RA, respectively. The solid diamonds and circles represent the probes not rejected by the HEIDI test. The yellow star indicates the top variant chr22:39252907. The second plot shows −log_10_(*P* values) of the variant’s associations for *SYNGR1*. The third plot shows −log_10_(*P* values) of the variant’s associations for chr22:39352274-39352775. The bottom plot (left) shows the importance of the peak-to-gene correlation identified by SCENIC+. The bottom plot (right) shows the potential regulatory relationships of TFs, chr22:39252907, chr22:39352274-39352775, and *SYNGR1.* (**D and E**) Violin plot illustrating the differences in chr22:39352274-39352775 accessibility (D), and *SYNGR1* expression (E) of Mature NK^dim^-FCGR3A cells across various genotypes at the chr22:39352907. (**F**) Effect sizes of variants from *SYNGR1* eQTL plotted against those for variants from the chr22:39352274-39352775 caQTL in Mature NK^dim^- FCGR3A cells. The orange dashed lines represent the estimate of *b*_xy_ at the top *cis*-caQTL. Error bars are the standard errors of variant effects. (**G**) Effect sizes of variants from RA GWAS plotted against those for variants from the *SYNGR1* eQTL in Mature NK^dim^-FCGR3A cells. The orange dashed lines represent the estimate of *b*_xy_ at the top *cis*-eQTL. Error bars are the standard errors of variant effects.

For example, rheumatoid arthritis (RA), an autoimmune disease, exhibited pleiotropic associations across nearly all cell types (Fig. 6B). Specifically, in Mature NK^dim^-FCGR3A cells, individuals carrying the A allele at chr22:39352907 showed increased chromatin accessibility at chr22:39352274-39352775 (*β* = 0.93), elevated *SYNGR1* expression (*β* = 1.22), and an increased susceptibility to RA (*β* = 0.14) (Fig. 6, C to G). Mutations in *SYNGR1* have been linked to various autoimmune diseases, and knockdown of *SYNGR1* expression in NK cells may affect cytokine secretion (*49*). Furthermore, we found additional evidence supporting the relationship between chr22:39352274-39352775 and *SYNGR1* within the GRN. TFs such as ASCL2, RUNX3, and TBX21 interact with binding sites within chr22:39352274-39352775, and peak-to-gene analysis suggests that this caPeak is linked to *SYNGR1* and may regulate its expression (Fig. 6C).

Additionally, in CD4 Tn-CCR7 cells, individuals carrying the T allele at chr17:39926578 exhibited decreased chromatin accessibility at chr17:39926349-39926850 (*β* = −1.32), reduced expression of *ORMDL3* (*β* = −1.43) and *GSDMB* (*β* = −1.31), and a decreased risk of developing asthma (β = −0.12) (fig. S6, A to D). The caQTL effect sizes of chr17:39926349-39926850 correlated positively with the eQTL effect sizes of *ORMDL3* and *GSDMB* and the GWAS effect size for asthma (fig. S6, C to D). The association between *ORMDL3*, *GSDMB*, and asthma is well-documented (*50*). *ORMDL3* encodes an endoplasmic reticulum-resident transmembrane protein that regulates sphingolipid biosynthesis by modulating the activity of serine palmitoyltransferase (SPT) (*51*). Overexpression of *ORMDL3* in CD4 T cells leads to SPT deficiency, reducing sphingolipid biosynthesis and attenuating TCR signaling. This may drive the differentiation of naive CD4 T cells toward a Th2 phenotype, thereby initiating the immune response associated with asthma (*52*). *GSDMB*, on the other hand, has been implicated in airway abnormalities related to asthma (*53*)

Further evidence from the GRN of CD4 Tn-CCR7 cells showed that TFs ETS1, KLF13, ZBTB14, and ZNF692 bind to motifs within chr17:39926349-39926850, targeting *ORMDL3* and *GSDMB*. This suggests that these TFs may regulate *ORMDL3* and *GSDMB* expression in CD4 Tn-CCR7 cells by interacting with binding sites within the caPeak, thus influencing the differentiation of these cells and contributing to asthma risk.

In conclusion, our findings reveal the potential regulatory effects of disease risk loci on chromatin accessibility and gene expression in immune cells, providing deeper insights into disease mechanisms.

### CIMA-CLM predicts chromatin accessibility and assesses the impact of noncoding variants in single cells

Large language models (LLMs), originally developed in the field of natural language processing (NLP), have shown significant potential in analyzing scRNA-seq data (*54*–*56*). However, integrating single-cell multi-omics data for downstream tasks remains a considerable challenge. To address this, we developed a novel cell language model, named CIMA-CLM, to predict chromatin accessibility and assess the effects of noncoding variants by taking as inputs chromatin sequences (501-bp in length) and single-cell gene expression data from our unique scATAC-seq and scRNA-seq datasets (Fig. 7A and fig. S7). The CIMA-CLM model is cell type-specific and begins with two parallel encoder branches. The first branch encodes input chromatin sequences into DNA embeddings using the pretrained HyenaDNA (*57*) model, while the second branch processes single-cell gene expression data to generate RNA embeddings using the pretrained scGPT model (*55*) (see Methods). The weights of these pretrained models are frozen with stop-gradient to speed up model training. To improve the model’s ability to generate more informative embeddings, two additional transformer-based encoders further process the DNA and RNA embeddings, respectively. CIMA-CLM incorporates a fusion decoder that integrates the DNA and RNA embeddings using multi-head cross-attention layers. This allows the model to capture relationships between chromatin sequences and gene expression data. Ultimately, the model predicts chromatin accessibility, measured by chromatin peaks, based on the fused features from both data types. To evaluate the model’s performance, we created a testing dataset for each of the 32 cell types by collecting chromatin sequences and single-cell expression data from 20 randomly selected individuals in our cohort. The remaining individuals’ data were used to construct training and validation datasets for model training (fig. S8A).

**Fig. 7.**
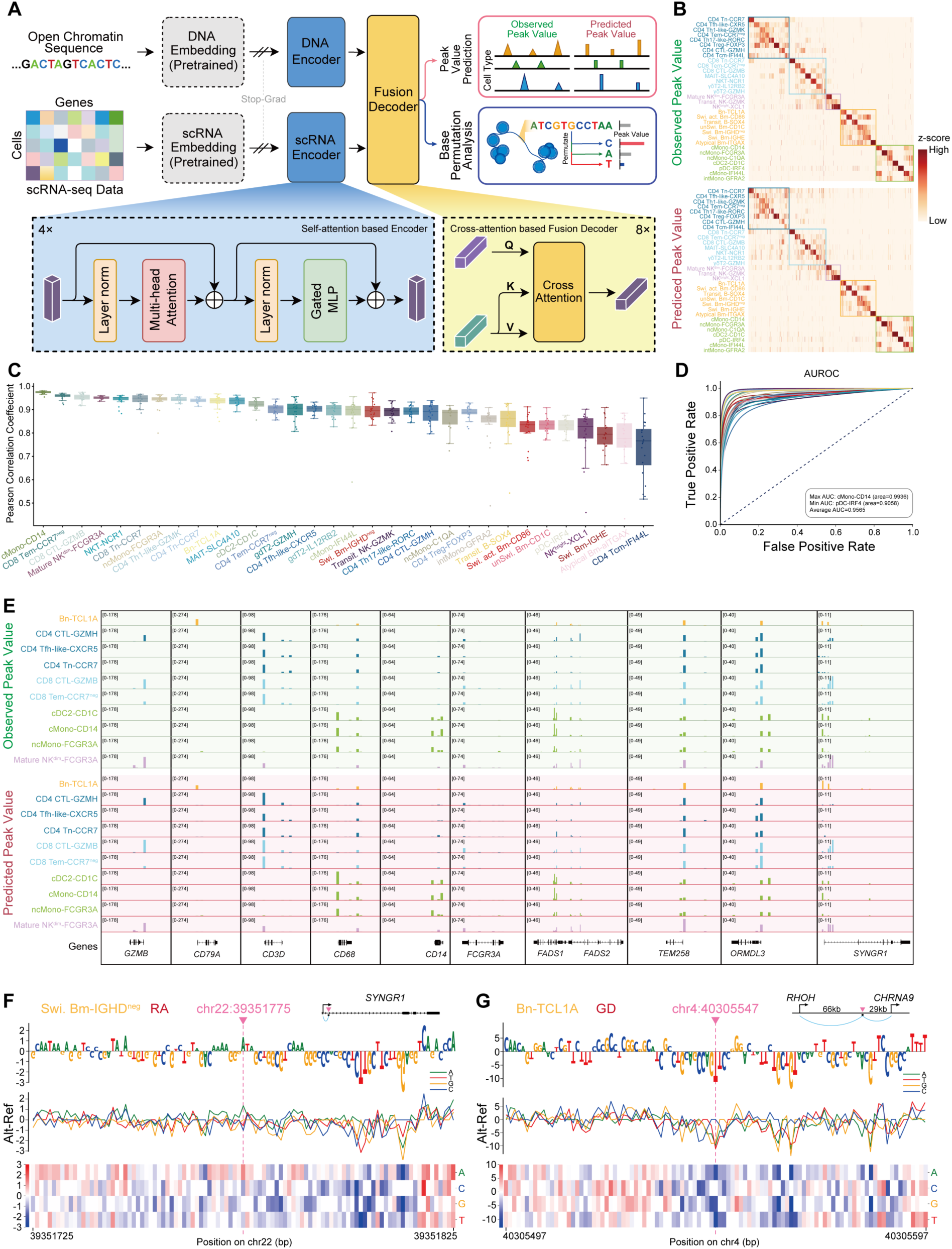
Prediction of chromatin accessibility and noncoding variation effect with single-cell multimodal language model CIMA-CLM. **(A)** Schematic diagram of the transformer-based CIMA-CLM model for predicting chromatin accessibility and assessing noncoding variation effects. (**B**) Heatmap showing the experimental peaks (top) and the predicted peaks (bottom) within the testing dataset based on the selected top 100 significantly differential chromatin open regions. Peak values are z-scored. (**C**) Chromatin accessibility prediction accuracy expressed as Pearson correlation coefficients for each individual per cell type within the testing dataset. (**D**) Area under the receiver operating characteristic curve (AUROC) for cell type-specific peak calling, comparing predicted peaks with real experimental signals. (**E**) Visualization of pseudobulk experimental observed peaks and predicted peaks for 10 cell types with their representative markers within the testing dataset. (**F and G**) In silico nucleotide mutagenesis effects on accessibility prediction in Swi. Bm-IGHD^neg^ cells for the chromatin sequence around the RA associated SNP variant at chr22:39351775 (F) and Bn-TCL1A cells around the GD related SNP chr4:40305547 (G). Larger signals indicate higher predicted accessibility for altered sequences, while lower signals correspond to decreased predicted accessibility. The logo diagram (top) shows each nucleotide substitutions that result in the most significant change in accessibility, along with the corresponding predicted signals. The line chart (middle) and heatmap (bottom) illustrate the change in accessibility at each nucleotide position before and after the mutation.

To demonstrate the utility of the CIMA-CLM model in profiling chromatin accessibility, we compared the predicted chromatin peaks with experimental signals using the testing dataset. The results showed a high level of consistency between the predicted and observed scATAC-seq profiles across the 32 cell types (Fig. 7B), indicating that CIMA-CLM accurately predicted open chromatin regions in these cells. Analysis of Pearson correlation coefficients (PCC) further confirmed the model’s high accuracy in predicting chromatin peaks for each individual across various cell types, with median *r* values ranging from 0.7661 to 0.9612 and a mean of 0.8951 (Fig. 7C, fig. S8C and table S13). Additionally, the PCC values for each cell type across all individuals in the testing dataset showed similarly strong correlations (fig. S8C). We also observed that PCC values tended to decrease as the number of single cells decreased (fig. S8B), suggesting that accurate profiling of cell type-specific chromatin accessibility may be significantly influenced by the number of captured single cells. The results of the area under the receiver operating characteristic curve (AUROC) analysis for different cell types further validated the model’s high accuracy in predicting chromatin peaks, with AUROC values ranging from 0.9058 to 0.9927 and a mean of 0.9560 (Fig. 7D and table S13). This performance is generally better than the most advanced deep-learning models for predicting chromatin accessibility signals (*58*). To further assess CIMA-CLM’s performance, we visualized pseudobulk predicted peaks, quantified as mean values across all individuals in the testing dataset, for 10 cell types with representative markers. The model not only replicated common chromatin signals across cell types (10 genes) but also captured cell type-specific predictions. For example, B cells, T cells, and myeloid cells exhibited distinct peak signals at *CD79A*, *CD3D*, *CD68*, and *CD14*, respectively (Fig. 7E). Moreover, cytotoxic cells, such as CTLs and NK cells, displayed specific peak signals at *GZMB* and *FCGR3A*. Additional genes, including *FADS2*, *TEM258*, *ORMDL3*, and *SYNGR1*, also demonstrated excellent predictive accuracy.

Leveraging the single-base embedding capability of our CIMA-CLM model, we conducted in silico mutagenesis on open chromatin regions to predict the regulatory effects of risk variants. By systematically mutating each nucleotide within the 501-bp open chromatin regions in silico, we assessed the impact on chromatin accessibility by comparing the predicted chromatin peaks between the reference and altered sequences (Fig. 7, F and G). Most of these simulated single-nucleotide mutations in the flanking regions had minimal effects on predicted accessibility, whereas numerous nucleotide substitutions near the variant locus resulted in either an increase or decrease in chromatin accessibility. In GWAS of East Asian populations, the variant at chr22:39351775 within the *SYNGR1* gene was associated with an increased risk of RA (*21*). Our model predicted that the nucleotide substitution (G > A) at the risk variant chr22:39351775 would enhance chromatin accessibility in the chr22:39351725-39351825 region (Fig. 7F), with negligible effects from other nearby variants. These computational predictions aligned well with the SMR results (table S12). In Swi. Bm-IGHD^neg^ cells, when the chr22:39351775 variant shifts from G to A, caQTL analysis indicated increased chromatin accessibility at chr22:39351525-39352026 (*β* = 1.18), while GWAS results confirmed an increased risk of RA (*β* = 0.14). Additionally, this risk variant showed pleiotropic associations with both caQTL and GWAS results (*β* = 0.12) (table S2). To further validate the predictive power of our model, we examined additional known risk variants. For example, in Bn-TCL1A cells, the risk variant (C > T) at chr4:40305547, located between the *RHOH* and *CHRNA9* genes and associated with Graves’ disease (GD), was predicted to cause a significant reduction in chromatin accessibility (Fig. 7G).

These findings highlight the potential of CIMA-CLM to elucidate the functional consequences of genetic variants on chromatin accessibility, providing crucial insights into the mechanistic basis of disease-associated genetic variations. Moreover, this approach serves as a powerful tool for prioritizing variants for experimental validation and enhancing our understanding of the regulatory landscape underlying complex traits and diseases.

## Discussion

The cellular and molecular components of blood are critical indicators of human physiological and pathological states. In this study, we conducted a comprehensive multi-omics analysis of 10,247,216 immune cells and 1,568 biomolecules in the peripheral blood of 428 individuals from the Chinese population. This allowed us to systematically characterize sex-zand age-related changes in these blood components, and to explore the effects of genetic variation at the biomolecular level, as well as its influence on immune cell gene expression and chromatin accessibility, shedding light on potential regulatory mechanisms.

We first investigated variations in immune characteristics and biomolecules within the population. Individual-level data included immune cell proportions, lipids, metabolites, and blood biochemical indicators. Initial correlations between these indicators and sex or age were identified through basic dimensionality reduction analysis. However, MOFA enabled the extraction of a more extensive set of multi-omics features associated with sex and age. Similarly, age-related factors were identified from gene expression and chromatin accessibility data in immune cells. Further analysis revealed that these age-related gene expression and chromatin accessibility patterns were cell type-specific and varied between individuals, suggesting that gene expression variability within the population is subject to complex regulation, likely influenced by both genetic and non-genetic factors.

To address the current gap in large-scale single-cell studies of cCREs, we identified 338,036 cCREs through scATAC-seq of immune cells. A comparison of these cCREs with those in the ENCODE database revealed that 51.7% of the newly discovered cCREs in our study were cell-type specific. These functional genomic elements can regulate gene expression by binding TFs. To further explore this, we constructed immune cell-type specific GRNs based on scRNA-seq and scATAC-seq data, allowing us to identify key TFs that regulate gene expression. These GRNs may encode cellular identity, guide immune cell development (*59*), and exhibit cell type-specific TF activity. Moreover, the activity of these TFs varied between populations and was correlated with sex and age, indicating that population-level variation in gene expression is likely influenced by the activity of TFs binding to open chromatin regions.

Furthermore, we examined the impact of genetic variation on immune characteristics and biomolecular levels in plasma. We calculated correlations between genetic variants and both gene expression levels and chromatin accessibility for each cell type, finding that most eQTLs and caQTLs exhibited cell type-specific effects. Similarly, we assessed the associations between genetic variants and lipid and metabolite levels. Notably, several loci previously reported to show selective effects in northern and southern Chinese populations were identified in our eQTL, caQTL, and in-house GWAS results, providing additional insights into the role of these loci in immune cells and their related phenotypes.

We then employed the SMR method to detect pleiotropic associations among eQTL, caQTL, and GWAS results. This revealed several regulatory loci that simultaneously affect chromatin accessibility, gene expression, and disease risk, suggesting potential regulatory pathways from genetic variation to chromatin structure, gene expression, and ultimately, GWAS traits. Importantly, our cell type-specific GRNs enhanced the interpretability of these pleiotropic associations.

Finally, we utilized a large language model to predict chromatin accessibility in specific cell types and simulate genetic variation perturbations, enabling us to observe changes in target chromatin accessibility peaks. This approach offers a novel framework for studying the effects of genetic variation on chromatin structure and gene expression.

In conclusion, our study presents the most comprehensive multi-omics dataset for peripheral blood immune cells in the Chinese adult population. Key contributions of this resource include: (i) Biomolecular indicators in each sample, encompassing plasma lipid and metabolite data, as well as blood biochemical markers and their relationships with sex and age. (ii) Single-cell gene expression (count matrices) and chromatin accessibility data, including scATAC-seq signal traces for various cell types and ENCODE-compatible regulatory elements (b-cCREs and scCREs). (iii) An extensively annotated marker list and differentially expressed genes (DEGs) for PBMC subpopulations, accompanied by a detailed annotation language model. (iv) Cell type-specific GRNs. (v) Statistical summary data for eQTLs and caQTLs in immune cells.

Due to space limitations, we were unable to conduct an in-depth analysis of each component in this publication. As a result, we have made the relevant data publicly available to facilitate further research by other investigators, allowing for comparative analysis and experimental validation of our findings.

In summary, our study fills a critical gap in single-cell multi-omics cohort studies for the Chinese population and provides valuable insights into the complex regulatory mechanisms of the immune system. This resource lays a solid foundation for future research and the development of personalized medicine.

## EXPERIMENTAL MODEL AND SUBJECT DETAILS

### Data reporting

We did not use statistical methods to predetermine the sample size.

### Human subjects

Human subjects for this study were obtained from the population of adults undergoing health examinations at BGI Pengcheng Clinic in Shenzhen (Guangdong Province, China) from November 2021 to November 2022. Recruitment and study design for this study were approved by the Institutional Review Board of BGI (BGI-IRB 21157-T3), and all subjects signed a written informed consent form. The final 428 subjects enrolled in this study ranged in age from 20 to 77. All subjects were Chinese, 95% of whom were Han Chinese. Subjects with active disease and pregnant women were excluded from recruitment. Blood biochemistry data from the physical examination and information from the questionnaire were collected.

### Methods

#### Blood sample collection and processing

Each volunteer provided 5-10 mL of peripheral blood collected with EDTA anticoagulation tubes. Peripheral blood samples were transported to the laboratory within 4 hours to complete PBMC/plasma separation and freezing. Plasma was first separated from whole blood by centrifugation, and after two centrifugations, plasma was stored at −80°C in a refrigerator for subsequent WGS, Metabolomic and Lipidomic assays; for the bottom cell layer, SepMate tubes (Stemcell) were used to ensure rapid and efficient separation of peripheral blood mononuclear cells (PBMCs), which were then subjected to a cooling gradient and finally stored in liquid nitrogen. After gradient cooling, the separated PBMCs were stored in liquid nitrogen and PBMCs with a viability rate greater than 80% were recovered for subsequent single-cell library construction and sequencing.

#### Generation of the Genotype Data from WGS

Cell-free DNA from plasma samples was used to generate genotype data. The extraction and library construction of cell-free DNA were performed using the MGISP-960 High-throughput Automated Sample Preparation System (MGI, ShenZhen, China). Specifically, 220 µL of plasma sample was used, and cell-free DNA was extracted using the MagPure Circulating DNA KF Kit (Magen). The MGIEasy Cell-free DNA Library Prep Set (MGI, 1000012700) was then used to construct the sequencing library. WGS was performed with a sequencing depth of 30X.

#### Target metabolite quantification by HM Meta Assays

Metabolite quantification was performed using the HM Meta Assay (BGI, Shenzhen, China), which measures 700 compounds across multiple chemical classes including bile acids, amino acids, fatty acids, carboxylic acids, hydroxyl acids, phenolic acids and indoles. Serial dilutions of metabolite standards were used to generate calibration curves. Aliquots (20 μL) of standards or plasma samples were mixed with 120 μL of internal standard solution (methanol diluted metabolite internal standards) in 96-well plates. After centrifugation, the supernatants were transferred to new 96-well plates for derivatization with 10 μL of freshly prepared reagents (200 mM 3-NPH in 75% aqueous methanol and 96 mM EDC-6% pyridine in methanol) based on a modified version of the method by Han et al. Derivatization was carried out at 30°C for 60 min. Samples were diluted with 400 μL ice-cold 50% methanol before the supernatants were transferred to new 96-well plates for LC-MS analysis. UPLC-MRM analysis was performed using a Waters ACQUITY UPLC coupled to a SCIEX QTRAP 6500 PLUS mass spectrometer with an ESI source controlled by Analyst software. Chromatographic separation was achieved on a Waters ACQUITY BEH C18 column (1.7 μm, 100 mm × 2.1 mm). The instrument was operated in both positive and negative ion modes.

#### Target lipid quantification by HM lipid assays

Lipid quantification was performed using the HM Lipid Assay (BGI, Shenzhen, China) following previous reported, which measures 1600 lipid molecules across multiple lipid subclasses including 18 lipid classes-sphingomyelin (SM), ceramide (Cer), cholesterol ester (CE), monoacylglycerol (MAG), diacylglycerol (DAG), triacylglycerol (TAG), lysophosphatidic acid (LPA), phosphotidic acid (PA), lysophosphatidylcholine (LPC), phosphatidylcholine (PC), lysophosphatidylethanolamine (LPE), phosphatidylethanolamine (PE), lysophosphatidylinositol (LPI), phosphatidylinositol (PI), lysophosphatidylglycerol (LPG), phosphatidylglycerol (PG), lysophosphatidylserine (LPS) and phosphatidylserine (PS). Serial dilutions of lipid standards were used to generate calibration curves. Aliquots (20 μL) of standards or plasma samples were mixed with 120 μL of internal standard solution (precooled isopropanol with SPLASH internal standards, avanti) in 96-well plates. After centrifugation, the supernatants (80 μL) were transferred to new 96-well plates for LC-MS analysis in negative mode, while 20 μL supernatants were diluted into 100 μL with methanol for LC-MS analysis in positive mode. UPLC-MS/MS analysis was performed using a Waters ACQUITY UPLC coupled to a SCIEX QTRAP 5500 PLUS mass spectrometer. Lipid species of SMs, CEs, Cer, TAGs, DAGs, and MAGs were quantified in positive mode, while Phospholipids and lysophospholipids were quantified in negative mode with scheduled MRM setting.

#### Single-cell RNA and ATAC library preparation and sequencing

PBMC suspensions were barcoded using the DNBelab C Series Single-Cell Library Preparation platform with DNBelab C Series High-throughput Single-cell RNA Library Preparation Set V2.0 (MGI, 940-000519-00) and DNBelab C Series High-throughput Single-cell ATAC Library Preparation Set V1.0 (MGI, 940-000793-00). Approximately 22,000 cells were loaded per reaction to capture more than 10,000 single cells for scRNA-seq. After droplet generation, emulsion breakage, bead collection, reverse transcription, and cDNA library construction, the scRNA libraries were sequenced on DNBSEQ-T20 and DNBSEQ-T1 sequencer at the China National GeneBank (Shenzhen, China). The sequencing strategy included 30 bp read length for cell barcode and UMI in Read 1, 100 bp read length for cDNA and a 10 bp sample index in Read 2. For the remaining PBMC suspensions, 100,000-300,000 cells were fixed using 0.01% formaldehyde, followed by cell membrane lysis, nuclear transposition, and subsequent library construction using the DNBelab C Series High-throughput Single-cell ATAC Library Preparation Set. The scATAC libraries were also sequenced on a DNBSEQ-T20 sequencer at the China National GeneBank (Shenzhen, China) with the following sequencing strategy: 70 bp read length for cell barcode and DNA insertion in Read 1, 50 bp read length for DNA insertion and 10 bp for sample index in Read 2.

#### Single-cell RNA-seq raw data preprocessing

The raw sequencing reads from the Oligo libraries sequenced by DNBSEQ-T1 and the cDNA libraries sequenced by DNBSEQ-T20 were first corrected and filtered using PISA (*60*) (version 0.12b). Reads were then aligned to the Homo sapiens (human) genome assembly GRCh38 reference genome using STAR (*61*) (version 2.7.2b), retaining only those reads mapped to exons. The EmptyDrops (*62*) method was used to select valid beads, and the cosine similarity between beads was calculated based on the types and numbers of oligos. Similar beads (potentially originating from the same cell) were then merged. Finally, the PISA Count module was utilized to generate a UMI count matrix for cells and genes.

#### Single-cell RNA-seq data analysis

The scRNA-seq data analysis workflow was constructed based on existing computational tools, including Scrublet (*63*) (version 0.2.3), Scanpy (*64*) (version 1.9.3), rapids-singlecell (*65*) (version 0.8.1) and COSG (*66*) (version 1.0.1). This pipeline was applied to all scRNA-seq samples from the CIMA cohort. The workflow consists of the following steps:

##### Doublet detected

We used Scrublet to identify doublets in each library. Scrublet simulates doublets based on the observed data and calculates a doublet score for each single cell using a k-nearest neighbor classifier. Briefly, we first created an AnnData object using the raw count data from each library. Next, we ran the Scrublet function with the following parameters: expected_doublet_rate=0.06, min_counts=3, min_cells=3, log_transform=True, min_gene_variability_pctl=85, and n_prin_comps=30. The “call_doublets” function was then applied with a threshold=0.2, and the doublet detection results were added to the metadata for further analysis.

##### Quality control at single cell level

We used Scanpy to perform several QC metrics for each cell, including total counts, total number of genes, percentage of mitochondrial gene counts, and percentage of ribosomal gene counts. Next, we removed the identified doublets and retained cells within the following QC criteria: 500 < total number of genes < 6000, 1000 < total counts < 25000, and percentage of mitochondrial gene counts < 10%. Additionally, genes expressed in fewer than three cells were excluded.

##### Batch effect removal, dimensionality reduction, cell clustering and reclustering

We utilized the Scanpy workflow for batch effect correction, dimensionality reduction, cell clustering, and reclustering of single-cell transcriptomics data. Initially, we identified the top 3,000 highly variable genes (HVGs) from the quality-controlled data using the “sc.pp.highly_variable_genes” function with the “seurat_v3” flavor, excluding mitochondrial genes, ribosomal genes, the *MALAT1* gene, and immunoglobulin genes except for *IGHD*, *IGHE*, and *IGHM*, resulting in 2,859 HVGs. The raw count matrix was preserved in “.layers[’count’]” as a backup for future analysis. Next, the data were normalized using “sc.pp.normalize_total” (with target_sum=1e4) and log-transformed with “sc.pp.log1p”. Only HVGs were scaled, and linear dimensionality reduction was performed using PCA with “sc.pp.scale” and “sc.tl.pca”, setting zero_center=False to reduce memory usage. Batch correction was applied using the Harmony algorithm by “sc.external.pp.harmony_integrate” function with batch=’individual’. Subsequently, we computed neighbors on the first 19 Harmony dimensions using “sc.pp.neighbors” and conducted UMAP dimensionality reduction using “sc.tl.umap” (with min_dist=0.3) and clustering analysis using “sc.tl.leiden” (with n_iterations=2). Differential expression analysis was performed using COSG. After completing these steps, we removed doublet clusters and isolated clearly segregated cells from the B, T & NK, and Myeloid groups for sub-cluster analysis to further identify distinct cell populations.

In the reclustering process for cell types, we started fresh with the raw count matrix extracted from the cell types “.layers[’count’]”. After log normalization, HVGs were recalculated using the “sc.pp.highly_variable_genes” function with the “seurat_v2” flavor (with batch_key=’sample’), selecting all HVGs that passed the default thresholds while excluding mitochondrial genes, ribosomal genes, pseudogenes, and long noncoding RNAs (lincRNAs). The HVGs were then scaled, and PCA was performed, then applied the Harmony algorithm for batch correction using the rapids-singlecell “rsc.tl.harmony_integrate” function (with batch=’experimental_date’). Dimensionality reduction and clustering were carried out using the same functions and parameters as described previously. During reclustering, doublet clusters were removed, and cells from other mixed subsets were extracted, with the remaining cells undergoing the subset reclustering workflow again. To further identify cell types within subsets, we extracted subsets based on specific cell markers and performed iterative reclustering. In total, 6,484,974 high-quality single-cell transcriptomes were obtained, which were categorized into five major subtypes: CD4 T cells (2,320,103 cells), CD8 T and unconventional T cells (2,340,449 cells), B cells (566,569 cells), myeloid cells and HSPCs (754,770 cells), ILC and NK cells (503,083 cells).

##### Manual annotation of circulating immune cells

For the 6,484,974 high-quality PBMCs quality-controlled in scRNA-seq data, we performed fine annotation based on key marker genes and differentially expressed genes between clusters. Table S1B summarized the positive and negative expression of key marker genes in different levels of immune cells, which were obtained from a priori knowledge, relevant published studies, and databases such as Cell marker2.0 (*67*), Celltypist (*68*), PanglaoDB (*69*), and other databases. We also combined the marker genes (table S1C) calculated by COSG (*66*) for each fine clustering to determine the annotation. The main objective of the first round of clustering was to distinguish between different immune cell lineages. The most numerous cells were *CD3D*-expressing T cells, which could be further categorized into CD4 T cells, CD8 T cells, and other unconventional T cells. Multiple clustering was performed until CD4 T cells and CD8 T cells were completely separated based on the expression of *CD4*, *CD8A*, and *CD8B*. Since other unconventional T cells (e.g., NKT, γδT, and MAIT) were closer to CD8 T cells in terms of clustering and function, they were assigned to the same L1, i.e., CD8 T & other unconventional T. So, five L1 levels of cells were identified, which were CD4 T, CD8 T & other unconventional T, B cells, myeloid cells, and ILC & NK cells. Cells at the L1 were subclustered several times and the resolution was continuously optimized to determine the best classification. The L2 and L3 were defined according to the developmental stage of the cells and the expression of key functional genes, which can be considered as “cell states” at different granularities. We focused on the functional genes specifically expressed in different clusters and finally defined 73 immune cell types with high expression of specific functional genes and labeled their specific or highly expressed genes at L4. Notably, we were able to detect a small number of *CD34*-expressing hematopoietic stem cells (HSPC-CD34) when subclustering L1 myeloid cells, due to the sufficiently large number of cells. Since all immune cells in peripheral blood are derived from myeloid hematopoietic stem cells, we classified HSPC-CD34 as a separate category. Finally, cell types in the L4 layer were used in the subsequent analysis.

##### Single-cell ATAC-seq data preprocessing

Briefly, raw sequencing reads were aligned to the GRCh38 genome using Chromap (*70*) (version 0.2.3-r407). To capture valid cells with sufficient fragments, barcode beads with fragment counts below the knee point were considered empty droplet and subsequently removed. To address the phenomenon of multiple beads in one droplet during the experiment, d2c (*71*) (version 1.4.7) was used to merge barcode beads based on the same start and end site of fragment distribution within the beads. After merging the beads, MACS2 (*72*) (version 2.2.7.1) was utilized to identify enriched genomic regions, also known as peaks, and ultimately generate a matrix of cells multiplied by peaks for each library.

##### Single-cell ATAC-seq data analysis

The scATAC-seq data analysis workflow was constructed based on existing computational tools, including scDblFinder (*73*) (version 1.12.0), Signac (*74*) (version 1.9.0), and SnapATAC2 (*32*) (version 2.5.1). This pipeline was applied to all scATAC-seq samples from the CIMA cohort. The workflow consists of the following steps:

##### Per-sample quality control and doublet removal at the single-cell level

To detect doublets, we employed the scDblFinder software. Briefly, we first created a SingleCellExperiment object using raw count data for each library. We then ran the “scDblFinder” function with the parameters: clusters=TRUE, aggregateFeatures=TRUE, nfeatures=25, and processing=’normFeatures’. The doublet detection results were subsequently added to the metadata for further analysis. For detecting low-quality outliers, we utilized the Median Absolute Deviation (MAD) method. Initially, we performed several QC metrics calculations for each cell in each library using Signac, including fragment count, TSS enrichment, nucleosome signal, blacklist ratio, and the FRiP. We then calculated the median and MAD of the metric values for each library. Values that deviated more than four MADs from the median were flagged as outliers. Finally, we removed cells identified as outliers and doublets from all libraries. Additionally, we excluded cells that did not meet the following thresholds: fragment count < 1000, TSS enrichment < 5, FRiP < 0.6, and TSS proportion < 0.3.

##### Sample-level quality control and filtering

To identify sample-level outliers in this scATAC-seq dataset, we implemented a method based on the D-statistic, which measures the median Spearman correlation of each sample library with all other sample libraries. First, we calculated the mean QC metrics for each library, including fragment count, TSS enrichment, nucleosome signal, blacklist ratio, Mitochondria proportion, TSS proportion, and the FRiP. For each library, we then calculated the Spearman correlation with every other library using these metrics. The median of these correlation values was assigned as the D-statistic for that library. Libraries with a D-statistic less than or equal to 0.2 were flagged as outliers. Additionally, we excluded libraries that did not meet the following thresholds: mean fragment count < 1000, mean TSS enrichment < 5, mean FRiP < 0.6, mean TSS proportion < 0.3, and the number of cells < 500.

##### Batch effect removal, dimensionality reduction, and cell clustering

Here, we utilized the SnapATAC2 workflow to perform batch effect removal, dimensionality reduction, and cell clustering. First, we imported the data using the “snap.pp.import_data” function. Next, we selected the top 200,000 features using the “snap.pp.select_features” function and performed spectral embedding with the “snap.tl.spectral” function using default parameters. We then applied the Harmony algorithm for batch correction using the “snap.pp.harmony” function with the parameters: batch=’individual’, max_iter_harmony=30, and key_added=’X_spectral_harmony’. Finally, we conducted UMAP dimensionality reduction using the “snap.tl.umap” function with the top 30 Harmony dimensions and performed cell clustering analysis using the leiden algorithm. To maintain consistency of samples across different omics, we also removed some sample libraries to facilitate downstream analysis. After these steps, we removed clusters dominated by batch effects, resulting in a scATAC-seq dataset of 3,762,242 cells. To further identify distinct cell clusters, we conducted a second-round clustering using SnapATAC2, employing the same functions and parameters as previously described.

##### Cell type annotation, peak calling, and peak-to-gene linkage identification

As described in section “Integration of scRNA-seq and scATAC-seq Data”, we utilized the scglue (*23*) (version 0.2.3) to facilitate cell type-level cluster annotation. Following GLUE integration, we used L4 annotation of scRNA-seq data to identify cell clusters that are consistent with those in the scATAC-seq data. The integrated data was then embedded into a common low-dimensional GLUE space.

For the identification of cell type-specific candidate *cis*-regulatory elements (cCREs), we performed peak calling using the “snap.tl.macs3” function with the parameters: groupby=’final_annotation’, n_jobs=96. After peak calling, we merged the identified peaks of each cell type to generate a union peak set using the “snap.tl.merge_peaks” function, based on the GRCh38 genome reference. Finally, we constructed a peak matrix, representing peaks per cell, for the scATAC-seq dataset using the “snap.pp.make_peak_matrix” function.

#### Integration of scRNA-seq and scATAC-seq Data

To integrate unpaired scRNA-seq and scATAC-seq data for single-cell multi-omics analysis, we employed scglue for the integration process. First, gene location information in scRNA-seq data was obtained from the GRCh38.gtf file using the “scglue.data.get_gene_annotation” function. Subsequently, we constructed a prior regulatory graph using the “scglue.genomics.rna_anchored_guidance_graph(rna,atac)” function and mapped HVGs information onto the scATAC-seq data. Dimensionality reduction of the peak matrix from the QC-filtered scATAC-seq data was performed using the LSI algorithm with the “scglue.data.lsi” function and the following parameters: n_components=100, n_iter=15, use_highly_variable=True. The raw peak count matrix and LSI dimension matrix from scATAC-seq were used as the ATAC omics input for the GLUE model, configured with the “scglue.models.configure_dataset” function with parameters: prob_model=’NB’, use_highly_variable=True, use_rep=’X_lsi’, and use_batch=’sample’. The scRNA-seq data was downsampled by 50% to reduce data volume and approximately align the number of cells with scATAC-seq. The original gene count matrix and PCA dimension matrix were used as the RNA omics input for the model, configured with the “scglue.models.configure_dataset” function using the parameters: prob_model=’NB’, use_highly_variable=True, use_layer=’counts’, use_rep=’X_pca’, and use_batch=’sample’.

To construct the model, we extracted the highly variable parts of both omics from the prior regulatory graph using the “guidance.subgraph” function to generate guidance_hvf. Model training was performed using the data from both omics along with guidance_hvf by utilizing the “scglue.models.fit_SCGLUE” function with the following parameters: init_kws={’h_dim’:512, ‘random_seed’:666} and fit_kws={’data_batch_size’:8192}. Upon completing the model training, we extracted the aligned X_glue matrices from both omics using the “glue.encode_data” function. Finally, cell type annotation from the scRNA-seq data was transferred to the scATAC-seq data through the aligned X_glue matrix using the “scglue.data.transfer_labels” function.

#### PCA analysis of bulk data

PCA analysis can directly reduce dimensionality and identify the main sources of variation in various biological datasets. In this study, we used metabolomics, lipidomics, blood biochemistry, scRNA-seq proportion, and scATAC-seq proportion bulk data as inputs for PCA analysis. Initially, the raw data were normalized to a 0-1 scale to ensure equal contribution from each feature. Notably, we removed features with missing values exceeding one-third in lipid data, and then filled the remaining features into the missing values with the averages. We also utilized the “sc.pp.highly_variable_genes” function to compute highly variable features. Then, PCA was conducted using the “sc.tl.pca” function in Scanpy (*64*) (version 4.0.3) with default parameters. The first two principal components were selected for visualizing sex and age distribution.

#### MOFA analysis of bulk data

We integrated data from 428 individuals, including 19 blood biochemical markers, 1,228 lipids, 321 metabolites, and proportions of 73 cell types identified by scRNA-seq and 65 cell types identified by scATAC-seq, as inputs for MOFA analysis. Using mofapy2 (*25*) (version 0.7.1), we trained the MOFA model by defining these datasets as five distinct “views” (representing different omics layers). In the first round of analysis, we applied a single-group MOFA+ framework to train the MOFA model. In the second round, we used a multi-group MOFA+ framework, setting sex batch as the “group”. In this framework, features were group-centered before model fitting, allowing us to identify shared sources of variability across groups and thereby mitigate group-specific effects. From the converged MOFA model, we extracted both the sample-factors matrix, which contained the values of the latent factors, and the factors-feature weight matrix using mofax (version 0.3.6). To identify factors associated with covariates such as age, sex, and BMI, we used the “cal_correlation” and “nonparametric_test_category” functions in scPAFA (*75*) (version 0.1.3). We then applied the “runumap_and_plot” function for UMAP dimensionality reduction and visualization of these factors. The factors-feature weight matrix can be further utilized to provide a biological interpretation of the factors.

To identify the MOFA factors most correlated with clinical indicators, we performed a correlation analysis between MOFA factors and clinical indicators across individuals, including age, sex, and BMI. Additionally, we counted the number of top-ranked features on Factor 2 for each view (five bulk data types).

#### MOFA analysis of single-cell gene expression and chromatin accessibility data

We log-normalized the single-cell gene expression matrix and then used the “get_pseudobulk” function from decoupler (*76*) (version 1.6.0) to convert it into pseudobulk gene expression matrices for each cell type. We retained only pseudobulk samples where the cell count exceeded 10, and the mean gene expression of all cells within each sample in each cell type was used as the value for the pseudobulk gene expression matrix. For each cell type, we retained only the top 2,000 features with the highest coefficient of variation across pseudobulk samples. For the single-cell chromatin accessibility matrix, we applied TF-IDF transformation using the muon (*77*) (version 0.1.6), followed by pseudobulk integration and feature selection in the same manner as for the single-cell gene expression matrix. Pseudobulk gene expression matrices and peak accessibility matrices were combined as input for MOFA. To integrate the pseudobulk RNA-seq and ATAC-seq results from different cell types, we applied a multi-group MOFA+ framework, setting the combination of cell types and omics (RNA and ATAC) as ‘view’ and batch as the ‘group’. From the converged MOFA model, we conducted the aforementioned downstream analyses, similar to those performed with the bulk MOFA model.

#### Variance partitioning analysis

To quantify the contribution of different sources to gene expression variance, we utilized the variancePartition (*28*) package (version 1.33.14) for our population-scale scRNA-seq data. We first selected cell types expressed in at least 10 cells per individual across more than 200 individuals. For each gene, we constructed a pseudobulk expression matrix for each cell type and individual. Lowly expressed genes and low-quality pseudobulk samples with fewer than 5,000 features were filtered out. Using variancePartition, we then analyzed all pseudobulk samples to determine the percentage of variation attributable to cell type, individual, and residual. To visualize the distribution of gene function in expression variation, we collected GO annotations from geneontology.org (*78*), MsigDB (*79*), and BIOCARTA (*80*). We also curated specific gene sets related to sex, aging, and housekeeping etc. For each gene set, we calculated the mean partitioned variance attributable to cell type, individual, and residual, which allowed us to compare the partitioned variance across different gene sets based on their associated gene variation percentages.

#### Construction of Gene Regulatory Networks

##### Metacell Calculation

We utilized SEACells (*81*) (version 0.3.3) and used “SEACells.core.SEACells” function to calculate metacells separately for scRNA-seq and scATAC-seq data at a ratio of approximately 50:1 for each cell type per sample, assigning SEACell_IDs. For cell types with fewer than 100 cells in a sample, all cells were assigned the same SEACell_ID. SEACell_IDs were stored in the metadata. The “SEACells.core.summarize_by_SEACell” function was then used to aggregate the raw counts of cells with the same SEACell_ID into metacells, retaining sample informations and cell type annotations. This process resulted in 142,389 scRNA-seq metacells and 86,366 scATAC-seq metacells.

##### Construction of Pseudo-Multi-Omics Metacell Data

To construct pseudo-multi-omics data, we randomly selected metacells from scRNA-seq and scATAC-seq for each cell type per sample based on the minimum number of metacells between the two omics, pairing them and assigning a common new metacell barcode. Cells not assigned a new metacell barcode were filtered out, resulting in 72,195 pseudo-multi-omic metacells.

##### Construction of pycisTopic Object and Topic Modeling

The scATAC-seq metacell data from pseudo-multi-omics data were preprocessed, and a pycisTopic object was constructed using the “create_cistopic_object” function from pycisTopic (*36*) (version 1.0a0). Topic modeling was performed on this pycisTopic object using the “run_cgs_models_mallet” function with the following parameters: n_topics=[5,30,50,80,100,130,150], n_cpu=96, n_iter=100, random_state=555, alpha=50, alpha_by_topic=True, eta=0.1, eta_by_topic=False, reuse_corpus=True. The number of topics was evaluated using the “evaluate_models” function, and a model with 130 topics was selected using the “add_LDA_model” function.

##### Candidate enhancer region inference and Motif Enrichment Analysis

To infer candidate enhancer regions, two methods were used: binarizing region-topic probabilities and calculating Differentially Accessibile Regions (DARs). The “binarize_topics” function from pycisTopic (using both ‘otsu’ and ‘ntop’ methods) was used to binarize the topics, selecting either all regions passing the threshold for each topic or the top 3,000 regions per topic. For data containing all cell types, the “impute_accessibility” function was used to estimate regional accessibility for the pycisTopic object, followed by scaling with the “normalize_scores” function (scale_factor=10^4). Highly variable regions were identified using the “find_highly_variable_features” function. DARs were computed using the “find_diff_features” function for per L3 cell type, per L4 cell type, sex and age groups within each cell type. The same analysis was performed for subsets data within each L4 cell type to identify DARs between L4 cell type. These regions will be inferred as candidate enhancers for next analysis.

Custom cistarget databases based on CIMA data were created using pycisTarget (*36*) (version 1.0.3), and motif enrichment analysis of candidate enhancers was conducted using the “run_pycistarget” function with parameters run_without_promoters=True and save_partial=True.

##### Construction of Enhancer-Based Gene Regulatory Networks

To construct e nhancer-based gene regulatory networks, we followed the SCENIC+ (*36*) (version 1.0.1) workflow. First, pseudogenes and long noncoding RNAs (lincRNAs) we re filtered from the scRNA-seq metacell data, retaining 21,944 genes. The filtered pseudo-multi-omics scRNA-seq metacell data, pycisTopic object from the completed topic modeling, and candidate enhancers from the motif enrichment analysis were combined into a SCENIC+ object using the “create_SCENICPLUS_ object” function. Gene regulatory network construction was performed with the “ run_scenicplus” function, using parameters such as variable=’cell_type_l4’, species=’hsapiens’, assembly=’hg38’, tf_file=’utoronto_human_tfs_v_1.01.txt’, biomart_h ost=’http://sep2019.archive.ensembl.org/’, upstream=[1000, 150000], and downstream=[1000, 150000].

##### Downstream Analysis of Gene Regulatory Networks

The SCENIC+ object w as filtered using the “apply_std_filtering_to_eRegulons” function to retain eRegulons with regions that are positively correlated with gene expression, resulting in 404 eRegulons. The filtered results underwent AUC enrichment analysis again using the “score_eRegulons” function. Dimensionality reduction of the filtered A UC matrix was conducted using the “run_eRegulons_tsne” function. To identify high-quality eRegulons, we used the “generate_pseudobulks” function to create pseudobulks by merging every 10 cells and performed correlation analysis on datasets containing all cell types and subsets using “TF_cistrome_correlation” fun ction. High-quality eRegulons were selected based on a correlation absolute value threshold greater than 0.8 for all cell types and greater than 0.6 for subtype data, resulting in a total of 237 high-quality eRegulons. The filtered SCENIC+ object was exported to Loom format using the “binarize_AUC” and “export_to_ loom” functions for future analysis. The peak-to-gene results were exported using the “export_to_UCSC_interact” function, and transcription factor correlations w ere calculated and clustered using the “correlation_heatmap” function to explore different transcription factor modules. Transcription factor specificity scores (RS S) for each cell type were calculated using the “regulon_specificity_scores” func tion.

#### Generation and analysis of genotypes

##### Preprocessing and genotyping

The WGS FASTQ files from 455 samples underwent quality control using fastp (*82*) (version 0.23.2). ZBOLT (*83*) (MegaBOLT version 2.3.0) was then employed to perform read mapping, position sorting, duplicate marking, base quality score recalibration (BQSR), and variant calling. Subsequently, we used GATK (*84*) (version 4) for joint calling and variant quality score recalibration (VQSR). The quality control process using PLINK (*85*) (version 1.9) involved several steps to ensure the dataset was appropriately filtered before imputation. Initially, biallelic variants were extracted, followed by applying a missing rate threshold of 0.03. A sex check was conducted, and the dataset was further filtered based on a minor allele frequency (MAF) threshold of 0.01 and a Hardy-Weinberg equilibrium (HWE) p-value threshold of 1e-6. Heterozygosity rates were assessed, with individuals having rates outside of ±4 standard deviations being excluded. Finally, a genetic relatedness filter with a threshold of 0.125 was applied, resulting in a final dataset of 443 individuals and 7,222,808 markers.

##### Imputation of genotyping data

We performed imputation using Beagle (*86*) (version 4.1) against the 1KGP high-coverage panel (*40*), which includes 3,202 samples and retained only the variants with a DR2 greater than 0.3. Post-imputation quality control was conducted using PLINK, involving the extraction of biallelic variants, application of a missing rate threshold of 0.03, an MAF threshold of 0.01, and a HWE p-value threshold of 1e-6. After assessing heterozygosity rates, individuals with rates outside of ± 4 standard deviations were excluded. The final dataset consists of 441 individuals and 8,766,136 variants.

##### Principal component analysis of genotyping data

After removing Long LD regions from the genome and performing LD pruning, the genotype data from this study was merged with the 1KGP High Coverage dataset (*40*), followed by PCA using PLINK. The values of PC1 and PC2 were used for visualization.

#### Linear model-based xQTL detection

The cell type-specific pseudobulk gene expression and peak accessibility matrices generated for MOFA analysis were also leveraged for xQTL detection. We employed TensorQTL (*87*) (version 1.0.8) to conduct linear model-based *cis*-eQTL and *cis*- caQTL analyses. Briefly, genes and peaks present in fewer than 90% of samples within each cell type were excluded. The remaining pseudobulk gene expression and peak accessibility matrices underwent inverse normal transformation across samples and were subsequently used as phenotype inputs in TensorQTL. The covariates included sex, age, the first two genotype principal components (PCs), and PEER factors. PEER factors were derived from the top 2,000 highly variable genes or peaks; for cell types with fewer than 2,000 genes, all available genes were included. For *cis*-eQTL analysis, we focused on variants located within 1 Mb upstream or downstream of the gene’s transcription start site (TSS), while for *cis*-caQTL analysis, we included variants within 1 Mb on either side of the peak midpoint. We used the “map_nominal” function to get the nominal *P* values of each variant-gene or variant-peak pairs. Then we used the “map_cis” function to perform 10,000 permutations and generates phenotype-level summary statistics with empirical *P* value, enabling calculation of genome-wide FDR (q-value). Genes and peaks with a q-value < 0.05 were annotated as eGenes and caPeaks, respectively. The variant most strongly associated with each eGene and caPeak, identified by the lowest nominal *P* values, was extracted and designated as the top xQTL.

#### Estimating Genetic Effect Correlations Across Cell Types

We applied *r_b_* method (*88*) to estimate genetic effect correlations across cell types with more than 500 eGenes or caPeaks. Each cell type was designated as both the reference cell type and the query cell type. We selected the top *cis*-xQTLs in a reference cell type, and estimated *r_b_* between reference cell type and query cell type using these variants.

#### Comparison of eQTLs in CIMA, OneK1K, and ImmuNexUT

The overlap of eGenes among CIMA, OneK1K (*42*), and ImmuNexUT (*8*) was visualized using Venn diagrams. The Pi1 statistic (*π*1) was employed to assess the replication rate of top *cis*-xQTLs from OneK1K and ImmuNexUT in CIMA across different cell types. Briefly, cell types from OneK1K and ImmuNexUT were designated as the reference cell types, while cell types from CIMA were assigned as the query cell types. To calculate the Pi1 statistic, nominal *P* values of eQTLs in the query cell types that overlapped with top *cis*-xQTLs in the reference cell types were extracted. The Pi0 value was computed using the “pi0est” function from the qvalue (version 2.36.0) package, and Pi1 was defined as 1 − Pi0 for each reference-query cell type pair.

#### In house GWAS analysis of lipids and metabolites

To conduct the GWAS analysis of lipids and metabolites, we retained lipids that were missing (NA) in less than 25% of the samples, resulting in a total of 623 lipids and 321 metabolites for analysis using the MLMA model in GCTA (*89*) (version 1.94.0). Given the correlations among some lipids and metabolites, the significance threshold was adjusted to 5e-8 divided by the effective number of independent traits or tests. For lipids, we estimated the effective number of independent traits using an eigendecomposition analysis as described by Chen et al (*90*), identifying 6 independent traits. Consequently, the significance threshold for lipid-associated variants was set at 8.3e-9 (5e-8/6). For metabolites, we determined the effective number of tests by calculating the number of principal components needed to explain at least 95% of the variance in the metabolome (*91*) which was found to be 151. Thus, the significance threshold for metabolite-associated variants was set at 3.3e-10 (5e-8/151).

#### SMR analysis

The xQTL summary statistics for eGenes and caPeaks with nominal *P* values were extracted and formatted as BESD files. Additionally, GWAS summary statistics for the 14 lipids and metabolites, as well as for 52 immune-related diseases, were utilized in the SMR (version 1.3.1) analysis (*20*). We employed SMR to detect three types of pleiotropic associations within each cell type: (1) variant-caPeak-eGene, (2) variant-caPeak-Trait, and (3) variant-eGene-Trait. The default threshold of 5e-8 was used to select the associated xQTLs as instrumental variables for the SMR test. In each cell type, we applied the Bonferroni method to correct *P*_smr_ in the SMR results and reject associations with a *P*_HEIDI_ < 0.01. Associations with a *P*_smr_ less than 0.05/number of tests and a *P*_HEIDI_ ≥ 0.01 were considered significant pleiotropic associations.

#### CIMA-CLM model development

##### Data pre-processing and split

To establish cell type-specific CIMA-CLM models utilizing multi-omics data from scATAC-seq and scRNA-seq, we first generated consensus DNA sequence for each sample using the bcftools consensus command (*92*) (version 1.9), which relies on the obtained variant call format (VCF) data and the GRCh38 reference genome. The consensus sequences were then lifted over to GRCh38 coordinates. Chain files necessary for the LiftOver (*93*) were created using transanno software (version 0.4.4). The transanno software employs “PAF files” as input, which we generated by aligning the consensus sequences with GRCh38 using minimap2 (*94*) (version 2.17, with the option “-cx asm5”). Subsequently, each scATAC-seq peak region was mapped onto its respective sample’s consensus DNA sequence. We then extracted DNA segments within these regions using bedtools (*95*) (version 2.30.0), yielding a total of ∼127 million segments.

For the raw scRNA-seq data analysis, we employed Scanpy (version 1.10.1) to filter out undesirable genes and cells, normalize gene expressions to a scale of 10,000 counts per cell, and perform log-transformation. We then identified and selected a total of 5,000 HVGs for model development.

For each cell type, we aggregated peaks within each of 338,036 open chromatin regions across all cells per sample from the scATAC-seq matrix to represent peak values. These aggregated peaks underwent TMM normalization followed by log-transformation. We then calculated the ratio of unobserved significant chromatin accessibility for each cell type. This ratio was defined as the proportion of open chromatin regions that did not exhibit peak signals, represented by a peak value of zero. Cell types with the ratio greater than 90% were excluded, resulting in a total of 32 out of 65 cell types being retained for the cell type-specific CIMA-CLM model training (fig. S8B).

We randomly selected 20 samples to construct the testing dataset, while the remaining DNA segments from the other samples were allocated for training the CIMA-CLM models. Of these, 99.95% (∼120 million segments) constituted the training dataset, while the remaining 0.05% (∼60,000 segments) served as the validation dataset for early stopping and hyperparameter tuning.

##### CIMA-CLM model architecture

Our CIMA-CLM comprises three primary modules: a DNA sequence encoder, a single-cell gene expression encoder, and a fusion decoder (Fig. 7A and fig. S7). Both the DNA sequence encoder and the single-cell gene expression encoder share a similar architecture. Each begins with a pretrained model to capture hidden state embeddings, followed by four stacked transformer-based encoder blocks. The fusion decoder integrates the embeddings from the DNA sequences encoder and the single-cell gene expression encoder using eight cross-attention-based blocks.

##### DNA sequence encoder

For a given annotated cell type, let {**s***_r,j_*, **X***_j_*, p*_r,j_*} be an example training instance, where **s***_r,j_* represents the DNA segment of the *r*-th peak region from sample *j*, with length *L*_S_. 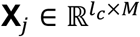 is the gene expression matrix with *l_c_* cells and *M* HVGs for the given cell type from the same sample *j*, and p*_r,j_* refers to aggregated peaked value of the *r*-th peak region from the sample *j*. For clarity and simplicity of notation, we denote the example instance as {**s***_i_*, **X***_i_*, p*_i_*} for the *i* -th example instance in the training dataset.

To encode the DNA segment, CIMA-CLM treated each nucleotide as a token, forming a DNA vocabulary that includes “A”, “T”, “C”, “G”, and “N” (a non-specific nucleotide), along with special tokens for padding, separation, and unknown characters. The DNA segment was subsequently tokenized, and padding tokens were added on the left side if necessary. The initial hidden state matrix, denoted as **E**_)_, was then encoded by adopting the pretrained HyenaDNA model with the model size of medium-450K (*57*):

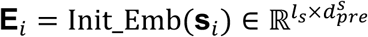

where *l*_s_ is the sequence length, 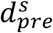 is the dimension of hidden state defined by the pretrained HyenaDNA model with the default value of 256. We chose HyenaDNA because it enables directly encoding DNA sequence at the single nucleotide level.

We further performed the shape transformation to obtain embedding matrix **S*_i_*** by adopting the adaptive average pooling to resize the length of sequence from *l_s_* to *l_seq_* and the gated MLP network to reshape hidden state dimension from *d_pre_* to *d_model_*:

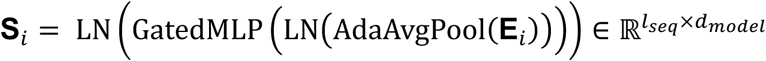

We set *l_seq_* and *d_model_* to be 501 and 512 in our experiment, respectively. LN represents LayerNorm layer. The Gated MLP network is given by GatedMLP(𝐙) = EGLU(𝐙𝐔_𝟏_ + 𝐛)K𝐔_𝟐_ + 𝐛, where 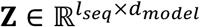, 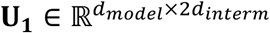, 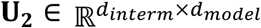, and *d_intern_* was set to 768 (fig. S7,left bottom). GLU represents the gated linear unit function, which is defined as GLU(𝐀) = 𝐀[:, : *d_model_*] ⊗ 𝜎(𝐀[:, *d_model_*]), where 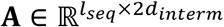 while 𝐀[:, :*d_model_*] and 𝐀[:, *d_model_*] represent the first and second half of matrix 𝐀, respectively. ⊗ refers to the Hadamard production, 𝜎() is the sigmoid function.

The embedding matrix of the input DNA segment **S***_i_* was then fed into a stack of four transformer-based encoder blocks to enhance the embedding capability. Each encoder block consists of a residual sub-block with an 8-head self-attention module, and a residual sub-block with a Gated MLP network. Both sub-blocks are preceded by a pre-LayerNorm layer. Each attention head separately transforms **S**_)_ into matrices of query **Q***_i_*, key **K***_i_*, and value **V***_i_*, where 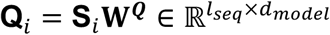, 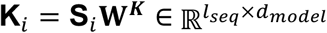, and 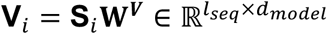. Queries represent the current information at each position and keys represent the information each position will be looking for to attend to. The output matrix was then produced as:

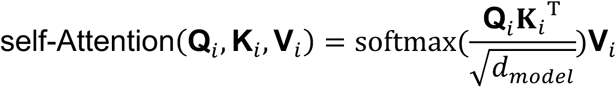

Subsequently, an 8-head self-attention mechanism was applied to capture joint attention across different representation subspaces at various positions. This process results in the representation denoted as **MHSA***_i_*:

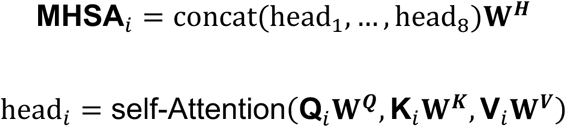

where 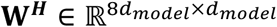 . The matrix **MHSA**_)_ was then input into the Gated MLP network, yielding the hidden state matrix 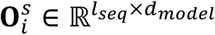 for the input DNA segment.

##### Single-cell gene expression encoder

The single-cell gene expression encoder has the same architecture as the DNA sequence encoder. Specifically, the single cell gene expression matrix **X***_i_* was first encoded by the scGPT model pretrained on 33 million normal human cells (*55*) (https://github.com/bowang-lab/scGPT), yielding a hidden state matrix 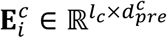, where *l_c_* is the sequence length, 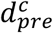 is the dimension of hidden state defined by the pretrained scGPT model with the default value of 512. The hidden state matrix then underwent the shape transformation to obtain embedding matrix 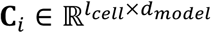, where *l_cell_* was set to 200 in our experiment. The embedding matrix 𝐂*_i_* was further encoded by a stack of four transformer-based encoder blocks, resulting in the hidden state embedding 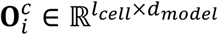 for the input single cell gene expression matrix.

##### Fusion decoder

The fusion decoder adopted cross attention mechanism to integrate the hidden state embeddings of 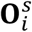 and 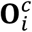 which carry information from DNA sequence and single cell gene expression, respectively. Specifically, the fusion decoder has a stack of eight blocks, and each block consists of a residual sub-block with an 8-head cross-attention module, and a residual sub-block with a Gated MLP network. Each attention head in the fusion decoder transforms the 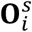 derived from DNA sequence into query 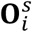 while the 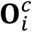 from single cell gene expression into key 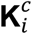 and value 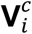, where 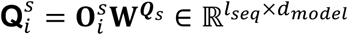, 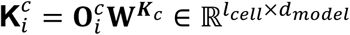, 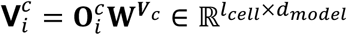. The cross-attention weight matrix was given by:

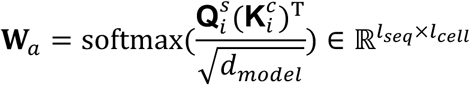

Then, the cross-attention output matrix was then yielded by:

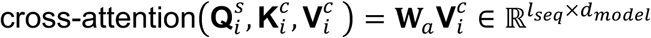

Subsequently, the output of the 8-head cross-attention, which was denoted as matrix **MHCA***_i_*, was achieved by:

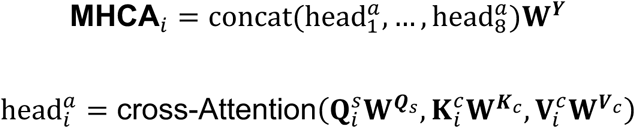

Where 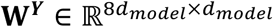. The matrix **MHCA**_)_ was then input into the Gated MLP network, yielding the final hidden state matrix 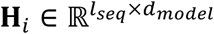.

We further transformed the shape of final hidden state matrix 𝐇*_i_* into the vector 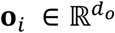 where we set *d_o_* to be 2048 in our experiment. The vector 𝐨*_i_* was finally fed into the output network to predict a peak:

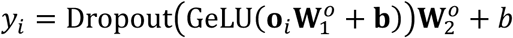

where 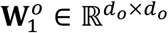, 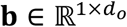, 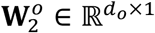, and *b* is a scalar bias for the final predicted peak value.

#### CIMA-CLM model training and evaluation

We used mean squared error (MSE) as our loss function during training phase:

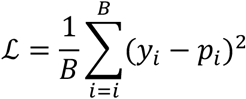

where 𝐵 is the batch size.

We developed and trained a separate CIMA-CLM model for each annotated cell type. The transformers framework (version 4.43.0) was employed to conduct the model training. We used AdamW optimizer (*96*) with the default parameters 𝛽_1_ = 0.9 and 𝛽_2_ = 0.999, and setting the weight decay to be 0.1. We set batch size of 72. We set initial learning rate to be 0.0006 and used cosine learning rate scheduler with 1,000 warmup steps. In general, these values for CIMA-CLM hyperparameters were determined in the way of combining grid search and manual tuning such that the optimal performance can be achieved. The maximum training step was 60,000, and the training process would stop early if the validation MSE value did not decrease in a total of 5,000 consecutive training steps. During the training of the CIMA-CLM model, we froze the weights of the pretrained components, HyenaDNA and scGPT. This resulted in the CIMA-CLM model having a total of 39 million trainable parameters.

For each cell type, we fully trained a CIMA-CLM model and used the testing dataset to evaluate the model’s performance. In addition to the calculation of MSE values, the model performance was assessed by the Pearson correlation coefficient (PCC) between the predicted peak values and the observed ones. Furthermore, we adopted Otsu function (*97*) to calculate thresholds that binarize the observed peak values in the test set. We then calculated the area under the receiver operating characteristic curve (AUROC) to evaluate the prediction accuracy of our CIMA-CLM model.

#### Single nucleotide perturbation analysis

For a given cell type, we extracted the sequence segments for each individual SNP site obtained in the testing dataset. We then performed simulated mutations on each nucleotide on those sequences, and used the CIMA-CLM model to predict the peaks of the mutated sequences. We adopted exponential function to rescale the predicted peak values as our model was trained based on the log-transformed peak values. For each individual at every SNP site, we calculated the difference in the predicted peak values between the mutated and original sequences, the mean value of those differences over the population in the testing dataset was used to perform perturbation analysis.

#### Model training environment settings

CIMA-CLM was implemented using Pytorch (version 2.0) and Python (version 3.9). All experiments were carried out on a cluster of four workstations. Each workstation was equipped with four NVIDIA A100 GPU addressing 40 GB RAM, and 128 Intel Xeon Gold CPUs addressing 250 GB RAM. It took approximately 18 hours to complete model training for each cell type-specific CIMA-CLM model.

## Acknowledgements

We would like to thank all our teams’ members and the China National GeneBank (CNGB) for their support.

## Funding

This work was supported by the National Key Research and Development Program of China (2022YFC3400400 to Xun Xu), National Science and Technology Innovation 2030 Major Program (2021ZD0200100 to Longqi Liu), China Postdoctoral Science Foundation (2023M732365 to Pengfei Cai, 2023M732369 to Yue Yuan), Boxiang Liu is supported by the Ministry of Education Singapore, under its Academic Research Fund Tier 1 (FY2023; 23-0434-A0001; 22-5800-A0001) and Tier 2 (MOE-T2EP30123-0015).

## Author contributions

Chuanyu Liu, X.X., L.L., J.Yin, Y.Zheng, and Wenwen Zhou conceived the idea; Chuanyu Liu, X.J., L.L., and X.X. supervised the work; Chuanyu Liu and J.Yin designed the experiment; Wenwen Zhou, Y.Zheng, Y.Y., S.D., and Y.Wang performed the majority of the experiments; Y.Zheng, Z.H., P.C., and Y.Y. performed data preprocessing and quality evaluation; Y.Zheng, Z.H., P.C., Y.B., S.Y., Y.Gao, Wenxi Zhang, Xinyu Zhang., and Y.Wei, analyzed the data; Y.H., T.Y., G.H., X.W., W.M., T.Q., and J.Yang provided technical support; Y.Liu, W.W., J.L., Z.Z., X.C., Xiru Zhang., Z.X., F.L., Y.Zhang, G.Z., S.H., Chang Liu, Y.L., B.W., Y.Li, Y.Gu, Wenwei Zhang, P.G., J.X., M.A.E., B.L., J.J. and J.W. gave the relevant advice; Wenwen Zhou, Y.Zheng, Z.H., P.C., Y.B., J.Yin, and Chuanyu Liu wrote the manuscript.

## Competing interests

The applications of this research are covered in pending patents. J.W. is founder of BGI-Shenzhen, and employee of BGI have stock holdings in BGI.

## Data availability

All data of this research have been deposited to CNGB Nucleotide Sequence Archive.

Visual exploration of processed scRNA-seq and scATAC-seq data is available in the XXXXX Database. All data were analyzed with standard programs and packages, as detailed above.

## Code availability

Additional information required to reanalyze the data reported in this paper is available from the lead contact upon request. Details of publicly available software used in the study are given in the Methods. All code is available from the corresponding author by reasonable request.

## Supplemental Materials

### Supplemental figures

**fig. S1.**
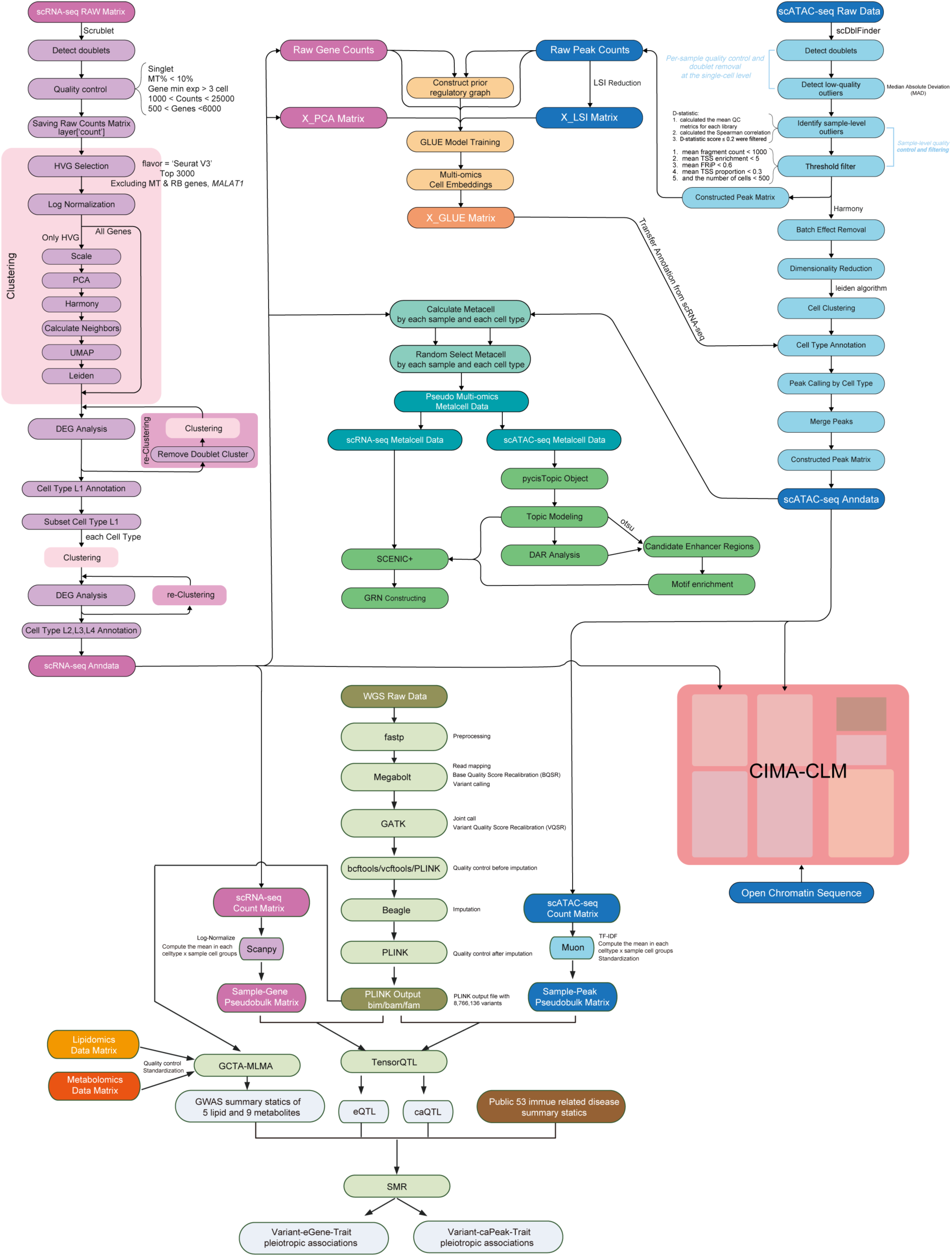
Comprehensive workflow for data processing and multi-omics integration analysis, related to Fig. 1 to 7. This flowchart illustrates the detailed data analysis pipeline of CIMA, including raw data processing from WGS, scRNA-seq and scATAC-seq, independent and integrated analysis of multi-omics data. The workflow includes the input and output files, software and algorithms used at each step.

**fig. S2.**
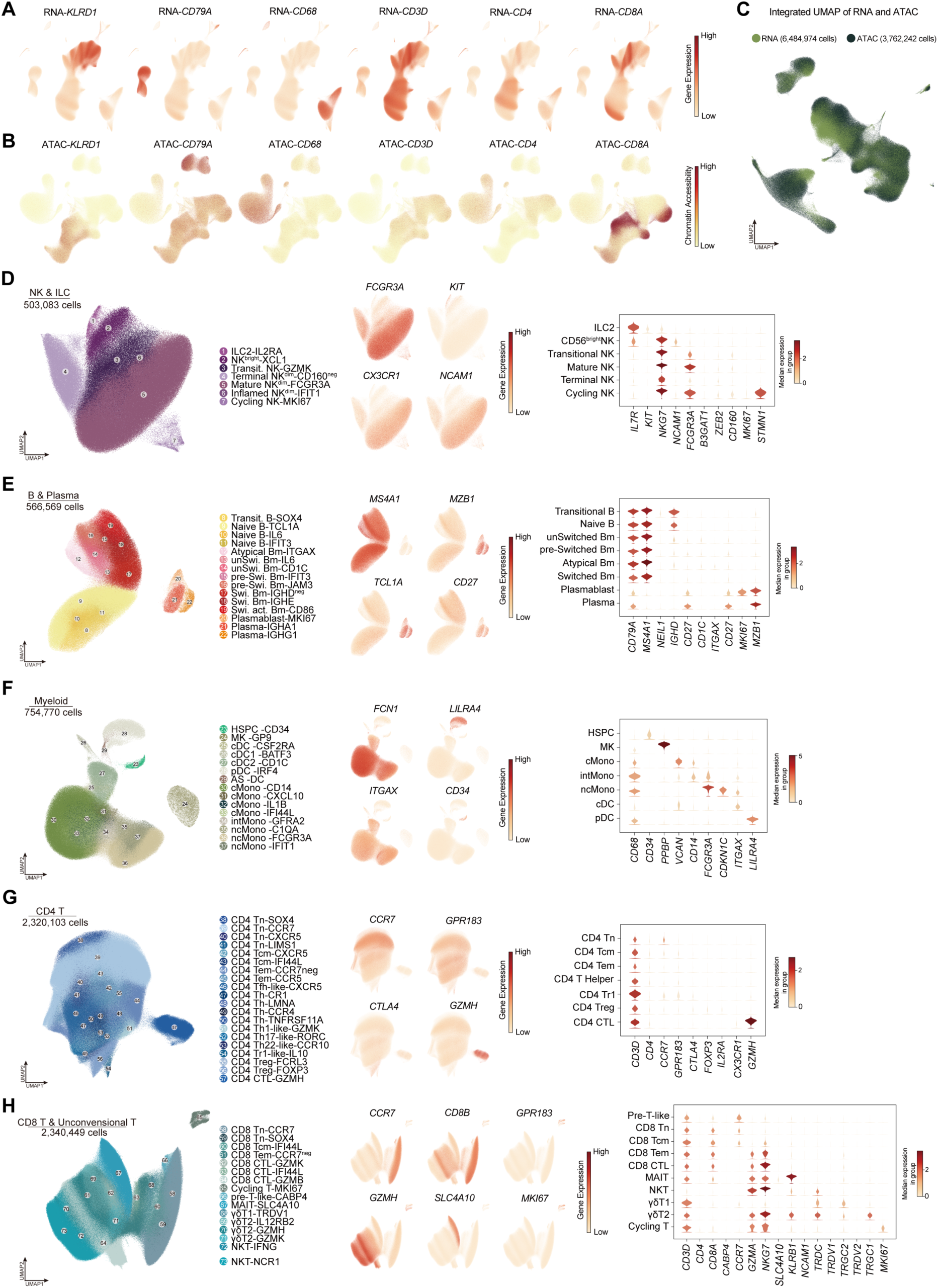
Classical marker genes for annotating L1-L3 immune cells, related to Fig.1. (**A and B**) UMAP plots showing the expression of classical marker genes for L1 immune lineages in scRNA-seq data (A), and gene activity of classical marker genes for L1 immune lineages predicted by scATAC-seq data (B). (C) The integrated UMAP plot showing the 6,484,974 cells captured by scRNA-seq and 3,762,242 cell captured by scATAC-seq. (**D**) Sub-clustering of 503,083 NK & ILC cells captured by scRNA-seq. The left UMAP plot showing the final annotation of L4 cell types. The middle UMAP plots displaying the expression of marker genes used to annotate L2 cells. The violin plots on the right showing the expression of marker genes used to annotate L3 cells. See Table S1 for marker genes for L4 cell types. (**E**) Sub-clustering of 566,569 B & plasma cells captured by scRNA-seq. The left UMAP plot showing the final annotation of L4 cell types. The middle UMAP plots displaying the expression of marker genes used to annotate L2 cells. The violin plots on the right showing the expression of marker genes used to annotate L3 cells. See Table S1 for marker genes for L4 cell types. (**F**) Sub-clustering of 754,770 Myeloid cells captured by scRNA-seq. The left UMAP plot showing the final annotation of L4 cell types. The middle UMAP plots displaying the expression of marker genes used to annotate L2 cells. The violin plots on the right showing the expression of marker genes used to annotate L3 cells. See Table S1 for marker genes for L4 cell types. (**G**) Sub-clustering of 2,320,103 CD4 T cells captured by scRNA-seq. The left UMAP plot showing the final annotation of L4 cell types. The middle UMAP plots displaying the expression of marker genes used to annotate L2 cells. The violin plots on the right showing the expression of marker genes used to annotate L3 cells. See Table S1 for marker genes for L4 cell types. (**H**) Sub-clustering of 2,340,449 CD8 T & uncoventional cells captured by scRNA-seq. The left UMAP plot showing the final annotation of L4 cell types. The middle UMAP plots displaying the expression of marker genes used to annotate L2 cells. The violin plots on the right showing the expression of marker genes used to annotate L3 cells. See Table S1 for marker genes for L4 cell types.

**fig. S3.**
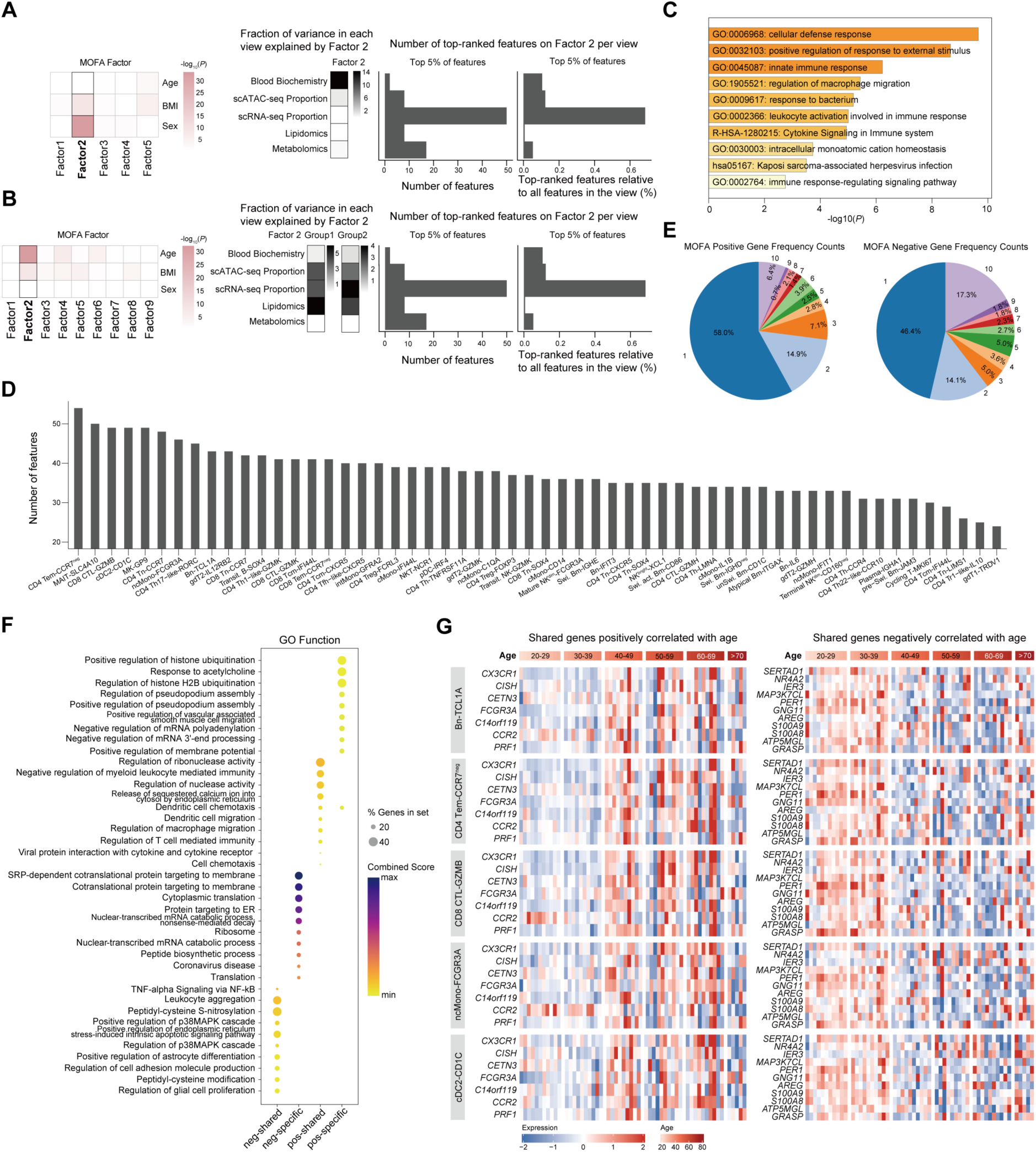
Comprehensive analysis of integrated cell types and genes in MOFA results, related to Fig. 2. (**A**) Overview of MOFA models using bulk multi-omics datasets, including cell type proportions from scRNA-seq, scATAC-seq, metabolomics, lipidomics, and blood biochemistry. Correlation analysis of MOFA factors with clinical indicators is shown for the raw model (left). Heatmap display the percentage of variance explained by each factor for each view (bulk data type) in the raw model (middle). Bar plots illustrate the total number of features and the relative amount of features (in respect to the number of view-specific features) among the top 5% highest-ranking features of factor 2 in the raw model (right). (**B**) Overview of MOFA models after correcting for sex-related effects using bulk multi-omics datasets in panel A. Correlation analysis of MOFA factors with clinical indicators is shown for the corrected model (left). Heatmap display the percentage of variance explained by each factor for each view (bulk data type) in the corrected model (middle). Bar plots illustrate the total number of features and the relative amount of features (in respect to the number of view-specific features) among the top 5% highest-ranking features of factor 2 in the corrected model (right). (**C**) GO function enrichment analysis of genes positively correlated with age (factor weight > 3) in cDC2-CD1C cells for factor 7 (RNA). (**D**) Bar plot showing the number of features passing the threshold (factor weight > 3 and factor weight < −3) for each view (cell type) in factor 7 (RNA). (**E**) Pie charts showing the fraction of gene frequency counts for genes positively correlated with age (factor weight > 3, left) and genes negatively correlated with age (factor weight < −3, right). (**F**) GO function enrichment analysis across four classes of genes: positive-specific, positive-shared, negative-specific, and negative-shared. (**G**) Normalized expression values of shared genes positively correlated with age (factor weight > 3, left) and shared genes negatively correlated with age (factor weight < −3, right) across different age stages in the Bn-TCL1A, CD4 Tem-CCR7^neg^, CD8 CTL-GZMB, ncMono-FCGR3A, and cDC2-CD1C cells.

**fig. S4.**
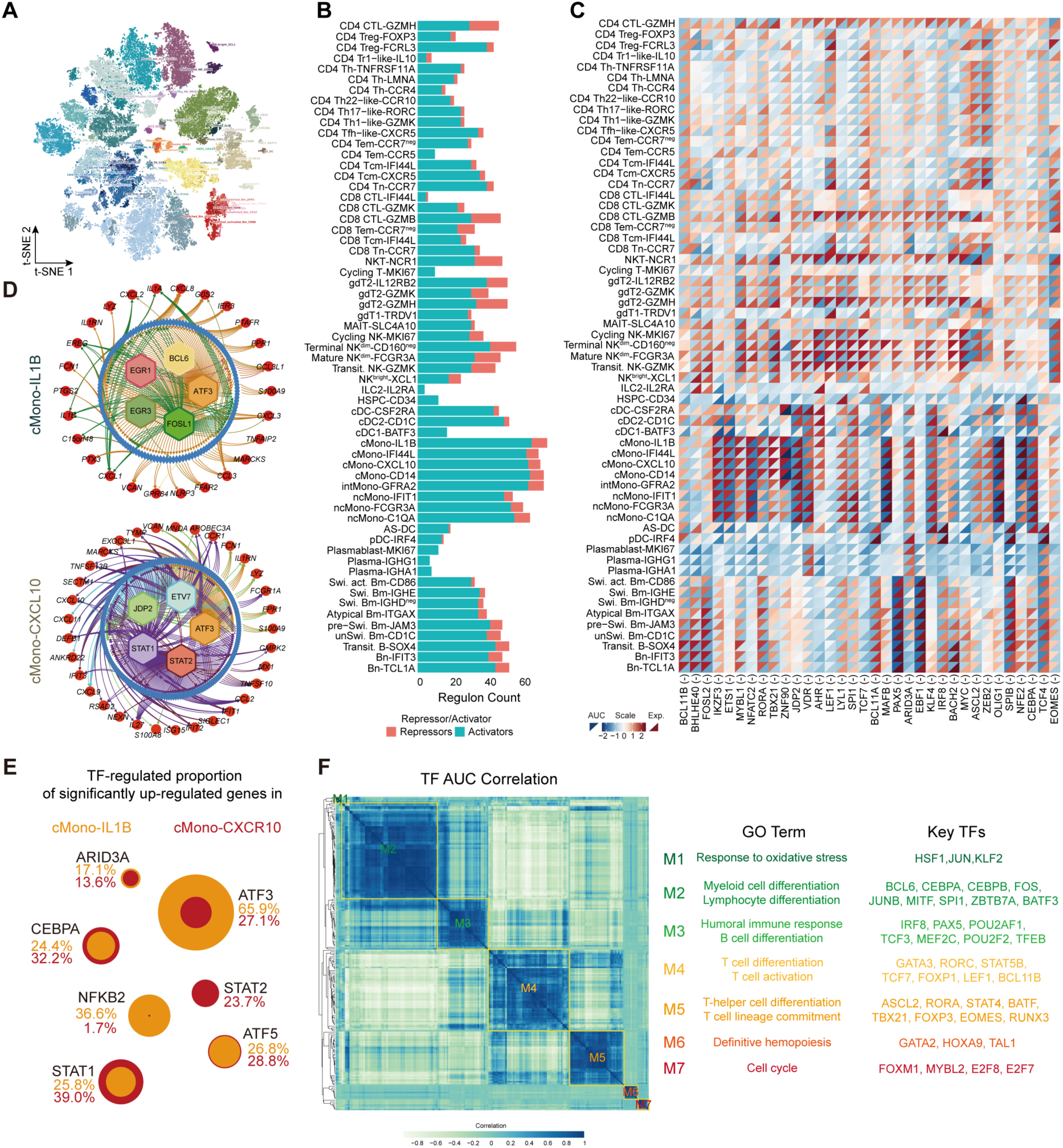
Construction of immune cell type-specific gene regulatory networks, related to Fig. 3. (**A**) t-SNE plot showing dimensionality reduction of cells based on eRegulon target gene and target region enrichment scores. (**B**) Bar plot showing the number of TFs with activity levels greater than 1 across different cell types. Activators are shown in green, while repressors are shown in red. (**C**) Heatmap showing the expression levels and activity of repressor TFs across different cell types. (**D**) Cell type-specific GRNs for cMono-IL1B (top) and cMono-CXCL10 cells (bottom). (**E**) Dot plot showing the proportion of upregulated genes regulated by TFs in cMono-IL1B (yellow) and cMono-CXCL10 cells (red). (**F**) Hierarchical clustering of TFs based on AUC correlations. Annotations of modules and representative TFs are shown.

**fig. S5.**
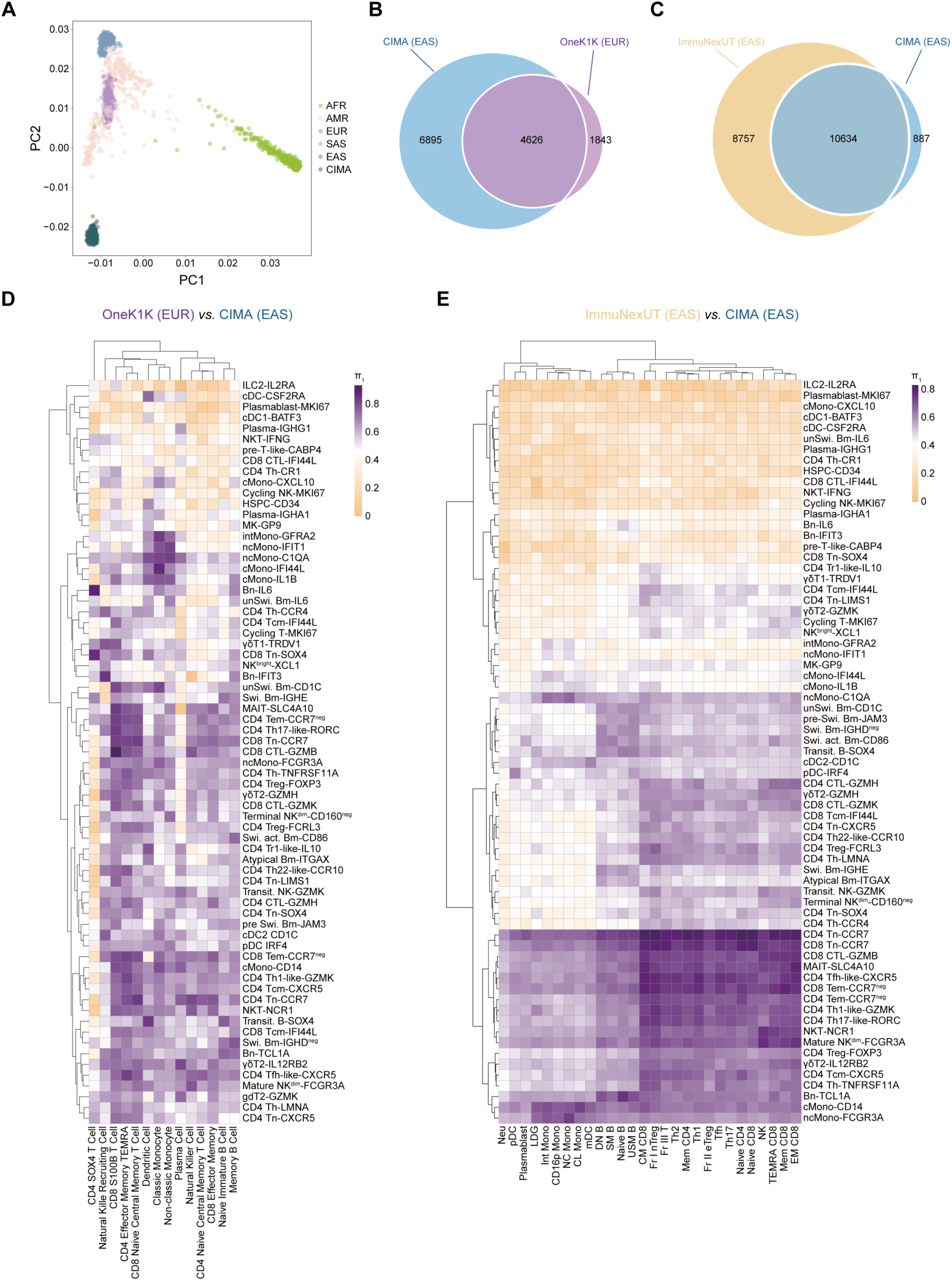
The comparison of xQTL in CIMA with the other datasets, related to Fig. 4. (**A**) PCA analysis shows the genetic relationship between the individuals from the CIMA and the 1KGP. Individuals from the CIMA are embedded with individuals with EAS ancestry. AFR, African; AMR, ad-mixed American; EAS, East Asian; EUR, European; SAS, South Asian. (**B and C**) Venn diagram showing the number of shared eGenes between CIMA and OneK1K (B), and between CIMA and ImmuNexUT (C). (**D and E**) Heatmap showing *π*1 statistic to quantitate the extent of eQTL sharing between each pair of cell type from OneK1K and CIMA (D), and from ImmuNexUT and CIMA (E). Rows represent reference eQTLs, while columns represent query eQTLs.

**fig. S6.**
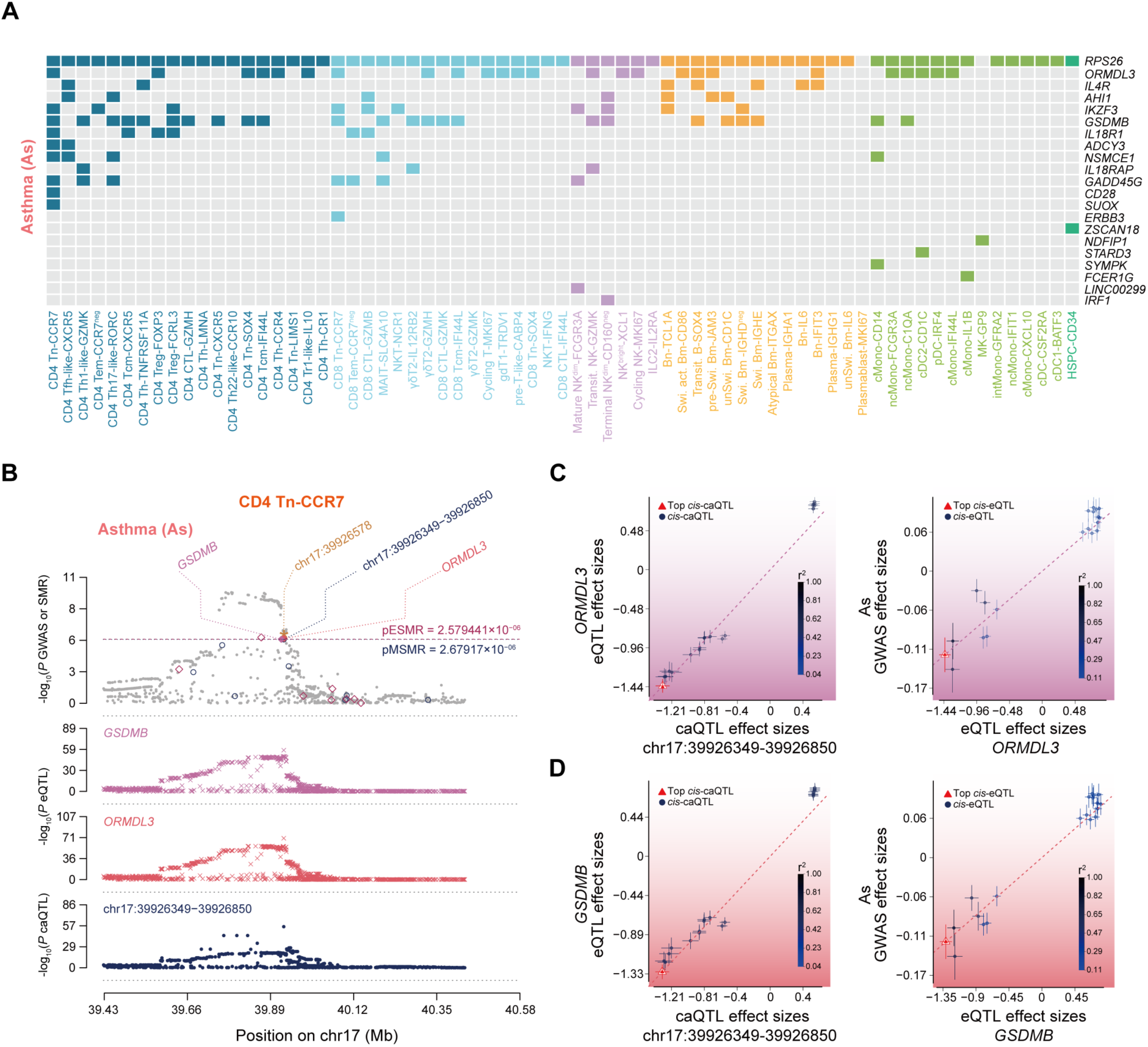
Genetic pleiotropic associations between xQTL and asthma, related to Fig. 6. (**A**) The eGenes detected in significant variant-eGene-disease SMR pleiotropic associations across cell types in As. (**B**) Results of variants and SMR associations across caQTL, eQTL, and As GWAS in CD4 Tn-CCR7 cells. The top plot shows − log_10_(*P* values) of variants from the GWAS of As. The red diamonds and blue circles represent −log_10_(*P* values) from the SMR tests for associations of eGenes and caPeaks with RA, respectively. The solid diamonds and circles represent the probes not rejected by the HEIDI test. The yellow star indicates the top variant chr22:39252907. The second plot shows −log_10_(*P* values) of the variant’s associations for *GSDMB*. The third plot shows −log_10_(*P* values) of the variant’s associations for *ORMDL3*. The bottom plot shows −log_10_(*P* values) of the variant’s associations for chr17:39926349-39926850. (**C**) Effect sizes of variants from *ORMDL3* eQTL plotted against those for variants from the chr17:39926349-39926850 caQTL in CD4 Tn-CCR7 cells (left). Effect sizes of variants from *ORMDL3* eQTL plotted against those for variants from the chr17:39926349-39926850 caQTL in CD4 Tn-CCR7 cells (left). Effect sizes of variants from As GWAS plotted against those for variants from the *ORMDL3* eQTL in CD4 Tn-CCR7 cells (right). The orange dashed lines represent the estimate of *b*_xy_ at the top *cis*-xQTL. Error bars are the standard errors of variant effects. (**D**) Effect sizes of variants from *GSDMB* eQTL plotted against those for variants from the chr17:39926349-39926850 caQTL in CD4 Tn-CCR7 cells (left). Effect sizes of variants from *GSDMB*. eQTL plotted against those for variants from the chr17:39926349-39926850 caQTL in CD4 Tn-CCR7 cells (left). Effect sizes of variants from As GWAS plotted against those for variants from the *GSDMB*. eQTL in CD4 Tn-CCR7 cells (right). The orange dashed lines represent the estimate of *b*_xy_ at the top *cis*-xQTL. Error bars are the standard errors of variant effects.

**fig. S7.**
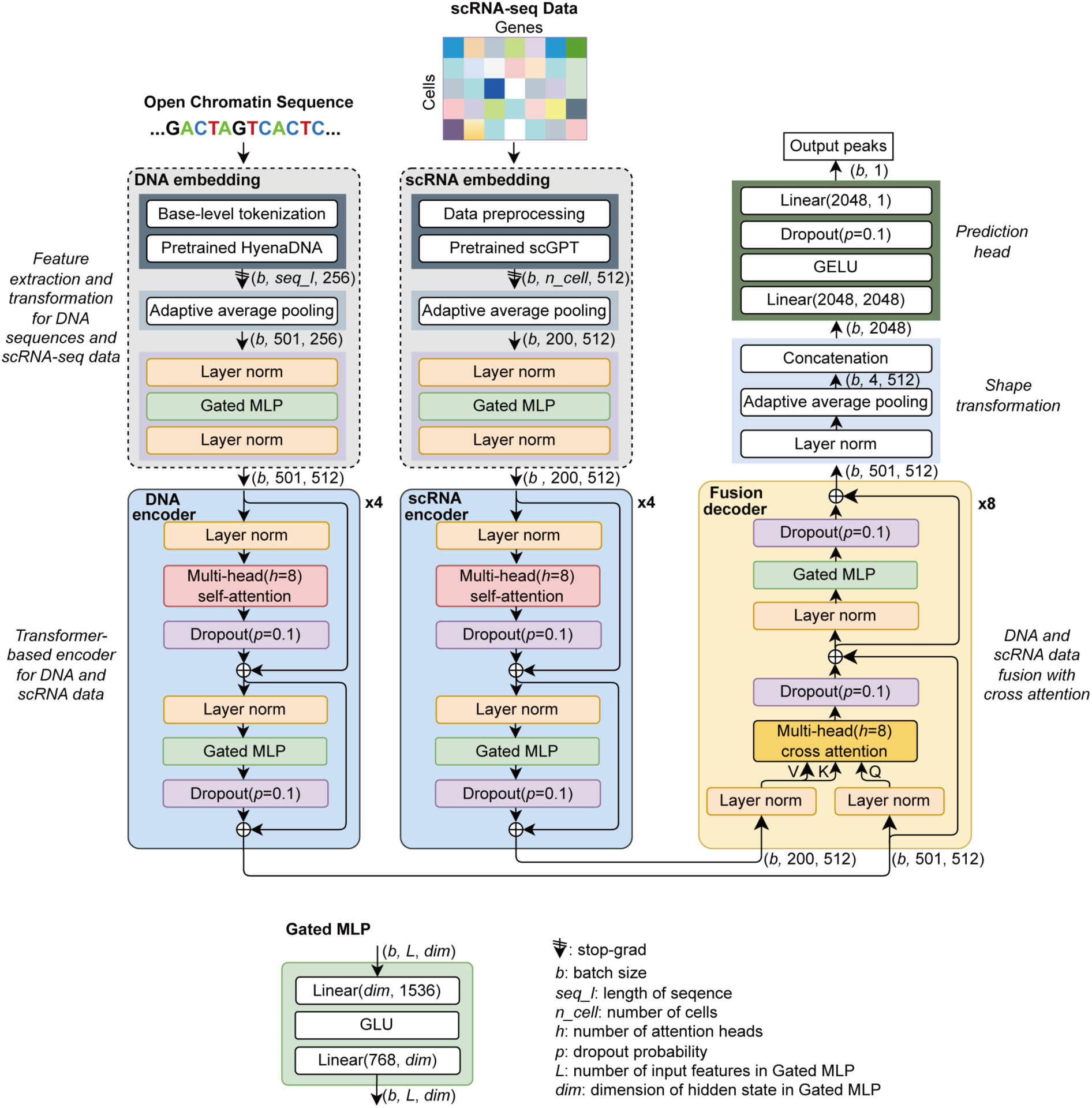
CIMA-CLM model architecture, related to Fig. 7. Output shapes are shown as tuples on the right side of the blocks. Definition of Gated MLP network blocks as a basic neural network layer was shown.

**fig. S8.**
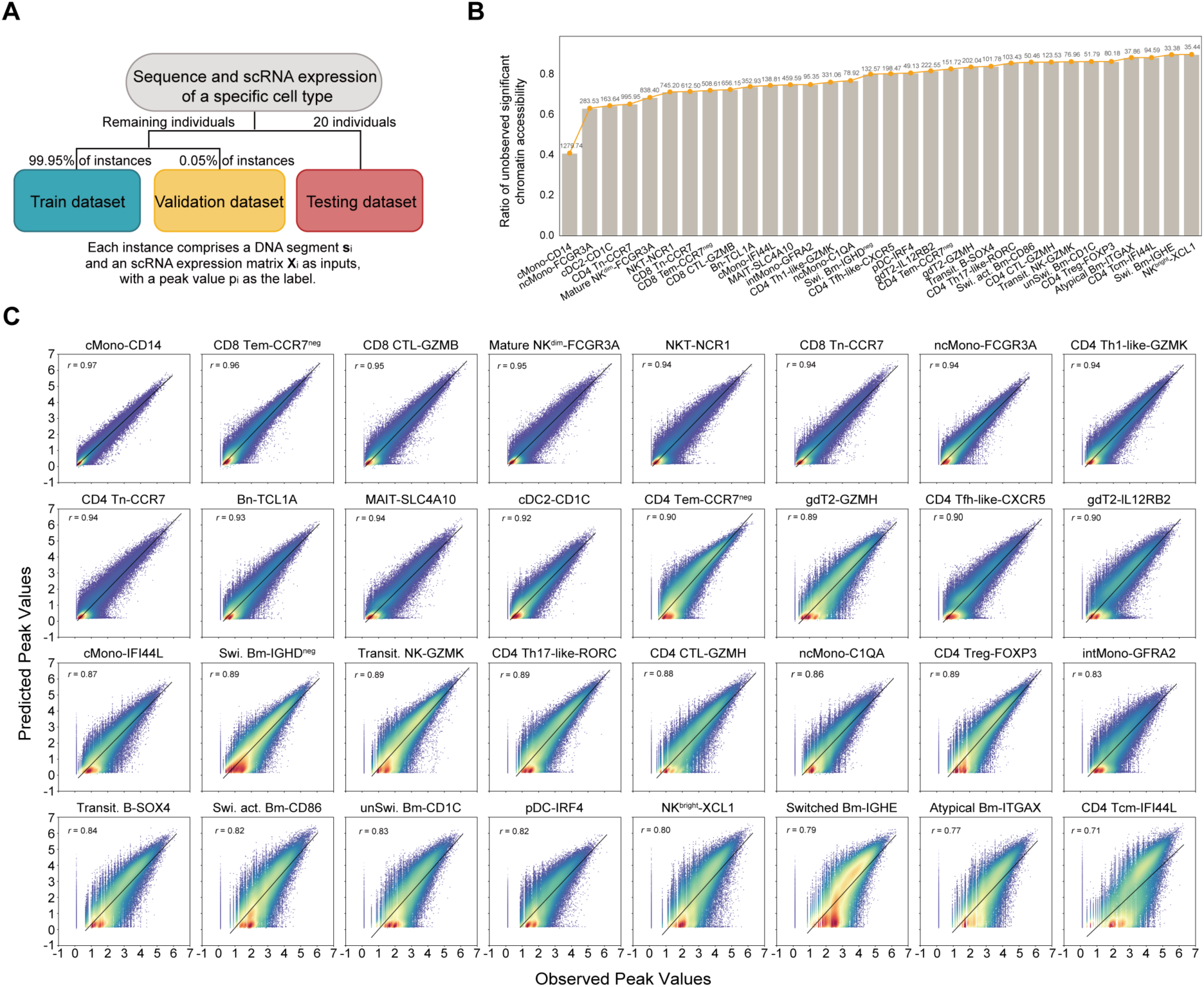
Data split for training and CIMA-CLM evaluation, related to Fig. 7. (**A**) Flowchart showing the creation of training dataset, validation dataset, and testing dataset for building cell type-specific CIMA-CLM model. Each example instance in the datasets is represented as {**s**_)_, **X**_)_, 𝑝_)_}^92@A^ (see Methods). (**B**) Bar plot showing 32 cell types with a ratio of unobserved significant chromatin accessibility less than 90%. The number above each bar represents the average number of single cells among for each cell type per individual in the whole dataset. (**C**) Scatter plot with Pearson *r* score showing the accuracy of predicting peaks among all individuals in the testing dataset for each cell type using CIMA-LM model.

### Supplemental tables

table S1. Metadata, multi-omics data, and immune cell type of CIMA, related to Fig. 1

table S2. Multi-omics molecular characteristics related to age and sex, related to Fig. 2

table S3. Peak accessibility of each circulating immune cell type detected by scATAC-seq, related to Fig. 3

table S4. Metadata of all eRegulons, related to Fig. 3

table S5. Immune cell type-specific eRegulons and TF activity, related to Fig. 3

table S6. caPeak with top variant, related to Fig. 4

table S7. eGene with top variant, related to Fig. 4

table S8. Selection loci under northern and southern China, related to Fig. 4

table S9. The catalog of 52 immune-related disease, related to Fig. 5

table S10. The significant loci associated with lipids and metabolites, related to Fig. 5

table S11. The significant SMR result of lipids and metabolites, related to Fig. 5

table S12. The significant SMR result of immune-related disease, related to Fig. 6

table S13. Performance of single-cell multimodal language model CIMA-LM, related to Fig. 7

